# Spike Count Analysis for MultiPlexing Inference (SCAMPI)

**DOI:** 10.1101/2024.09.14.613077

**Authors:** Yunran Chen, Jennifer M Groh, Surya T Tokdar

## Abstract

Understanding how neurons encode multiple simultaneous stimuli is a fundamental question in neuroscience. We have previously introduced a novel theory of stochastic encoding patterns wherein a neuron’s spiking activity dynamically switches among its constituent single-stimulus activity patterns when presented with multiple stimuli (Groh et al., 2024). Here, we present an enhanced, comprehensive statistical testing framework for such “**multiplexing**” or “**code juggling**”. Our new approach evaluates whether dual-stimulus responses can be accounted for as mixtures of Poissons either anchored to or bounded by single-stimulus benchmarks.

Our enhanced framework improves upon previous methods in two key ways. First, it introduces a stronger set of foils for multiplexing, including an “overreaching” category that captures overdispersed activity patterns unrelated to the single-stimulus benchmarks, reducing false detection of multiplexing/code-juggling. Second, it detects faster fluctuations - i.e. at sub-trial timescales - that would have been overlooked before. We utilize a Bayesian inference framework, considering the hypothesis with the highest posterior probability as the winner, and employ predictive recursion marginal likelihood method for the involving nonparametric density estimation.

Reanalysis of previous findings confirms the general observation of “code juggling” and indicates that such juggling may well occur on faster timescales than previously suggested. We further confirm that juggling is more prevalent in (a) the inferotemporal face patch system for combinations of face stimuli than for faces and non-face objects; and (b) the primary visual cortex for distinct vs fused objects.

## 1 Introduction

A central postulate in computational neuroscience holds that neurons encode sensory signals in a *repeatable* way. The activity elicited by a given stimulus is assumed to follow a consistent profile across time and across repeated presentations of that stimulus, with any variation falling within a given stable distribution. While repeatability may indeed occur when only one stimulus is present (e.g. in a typical, highly controlled laboratory setting with a single visual image or sound at a time), we recently documented evidence that when two stimuli are present, a subset of neurons show evidence of stochastic code juggling where the neuron’s spiking activity appears to dynamically switch between the activity patterns typically observed when each stimulus is presented alone (Caruso et al., 2018; Mohl et al., 2020; Glynn et al., 2019; Jun et al., 2022; Schmehl et al., 2024; Groh et al., 2024). Such stochastic juggling, also termed “Multiplexing”, is an exciting possibility because it could be information preserving: by leveraging the temporal dimension and/or by allowing different neurons to “switch” separately of one another, code switching may underlie our ability to perceive the individual elements of a crowded sensory scene.

This discovery was made possible by novel statistical tools which probed spiking activity of individual neurons aggregated by “trial” – one uninterrupted presentation of a sensory scene lasting several hundred milliseconds – with the spike train recorded within each trial summarized into a single *trialwise spike count*. The main statistical analysis framework was put forward in Caruso et al. (2018) who considered the hypothesis of **fluctuation across trials**: *when two stimuli A and B are presented together an individual neuron may encode stimulus A in some trials and stimulus B in the remaining trials – switching randomly between these options*. If neurons indeed fluctuate across repeated stimulus exposures, the double-stimuli trialwise spike count distribution must resemble a weighted mixture of the corresponding single stimulus distributions. Such a resemblance should be statistically testable with trialwise spike count data collected from three types of trials: stimulus A presented alone, stimulus B presented alone, or stimuli A and B presented together. Caruso et al. (2018) developed such a test under the additional assumption that the single stimulus benchmark distributions are each a Poisson. The Poisson assumption reinforces the concept of repeatability across single stimulus trials and is helpful when the A and B benchmark distributions are partially overlapping and hence, even when neural responses are fluctuating across repeated exposures, the AB distribution may not be visibly bimodal.

However, the existing statistical tools for detecting multiplexing have gaps that may contribute to both false positive and false negative discoveries. On the one hand, a chief limitation of the fluctuating-across-trials hypothesis is its inability to capture the notion that a neuron may switch one or more times during the course of a trial. If such sub-trial switching happens with any regularity, the corresponding trialwise spike count distribution could concentrate within an interval sandwiched between the A and B benchmarks, rather than splitting and spreading itself across the two. Accordingly, testing for across-trials fluctuations does not help with discovering existence of code juggling when it happens at a faster timescale.

On the other hand, the statistical test we developed in Caruso et al. (2018) is susceptible to falsely concluding in favor of cross-trial-fluctuations in situations where the double-stimuli spike count distribution is a weighted mixture of something other than the two benchmark Poisson distributions. For example, if the spike count distributions under A and B are, respectively, Poisson(50) and Poisson(80), and the same under AB is a 60-40 mix between Poisson(65) and Poisson(95), and data consists of 30 trials under each of the three conditions, the test would conclude in favor of across-trial-fluctuations on average four out of five times, even though the switching patterns on the AB trials are not an exact match to the benchmarks established on the A and B trials. This type of pattern is consistent with fluctuating activity, but the failure to coincide with the benchmarks makes it uncertain what information is contained in the signal.

Here, we fill in these gaps by refining the statistical testing framework to evaluate switching at either whole-trial or sub-trial timescale from trialwise spike count data. Furthermore, we expand the framework to incorporate consideration of generalized “overdispersion”– extra trial-to-trial variability of trialwise spike counts above and beyond what one should observe under the benchmarked-repeatability assumption. Our new statistical evaluation method, which we call Spike Count Analysis for MultiPlexing Inference (SCAMPI), retains the Poisson benchmark assumption for the two single-stimulus trial-wise spike count distributions. For double-stimuli exposures, we retain the **fluctuating-across-trials** hypothesis which entails that the corresponding trialwise spike count distribution must be a mixture of the two benchmark Poisson distributions. We refer to this possibility as **slow juggling**. To such slow juggling, we add the hypothesis of **fast juggling**, or **fluctuation at sub-trial timescales**: *when two stimuli A and B are presented together an individual neuron may encode stimulus A for some parts of a trial and stimulus B in the remaining parts – switching randomly between its choices within and across trials*. The “across trials” qualifier of this statement is important – in one trial the neuron may encode stimulus A 60% of the time, whereas in the next it may do so only 50% of the duration. Distinct from the slow juggling hypothesis, if the fast juggling hypothesis was true, the double-stimuli spike count distribution must equal a weighted mixture of *several* Poisson distributions each with a mean firing rate sandwiched between the benchmark A and B mean rates. Such distributional shapes can be statistically detected and formally tested from trialwise spike count data. (Although, the information is too coarse to discern the patterns of subtrial switching dynamic – such as when switches occur or what proportion of the time is spent responding to A vs B.)

In our new framework, slow and fast juggling together make up the multiplexing landscape where code juggling happens (see Figure 1). Both hypotheses describe the double-stimuli trialwise spike count distribution as a mixture of Poisson distributions. For slow juggling, it is a two-component mixture where the component means are pinned down at the benchmark mean firing rates. For fast juggling, the mixture may include several Poisson distributions but each of them must have their mean bracketed by the (means of the) benchmarks. As controls to slow and fast juggling, we consider two types of alternatives: one where the double-stimuli trialwise spike count distribution is a single Poisson satisfying the repeatability assumption, and the second where the double-stimuli spike count distribution is an overreaching mixture of Poisson distributions with some component mean firing rates falling outside the range defined by the two benchmarks. Clearly, if the AB distribution was Poisson(95), it would fall under the alternative model of a single-Poisson. Interestingly, so would the case where the AB distribution was Poisson(65). In theory, fast sub-trial fluctuation could produce a trialwise spike count distribution which is a single Poisson with mean between the benchmark values, but for it to happen, the fluctuation pattern has to be *repeatable* across trials – on each trial the neuron must spend exactly the same fraction of time coding for stimulus A. But a single Poisson distribution could also manifest without any switching, because of repeatable response by the neuron around a new mean firing rate that is between its rates for the two single stimulus cases. This has been hypothesized to occur as a normalization operation in which neurons (somehow) average their inputs (see Groh et al., 2024, for more discussion). By choosing single-Poisson as an “alternative” we allow for this alternative explanation. The second control scenario includes the example mentioned earlier, where A, B, and AB spike count distributions are, respectively, Poisson(50), Poisson(80), and a 60-40 mix between Poisson(65) and Poisson(95). Even though this situation can be seen as an example of switching (and such switching is potentially interesting), it is not consistent with the more precisely defined slow nor fast juggling, since the second mixture component mean lies outside the range defined by the benchmarks.

**Figure 1:**
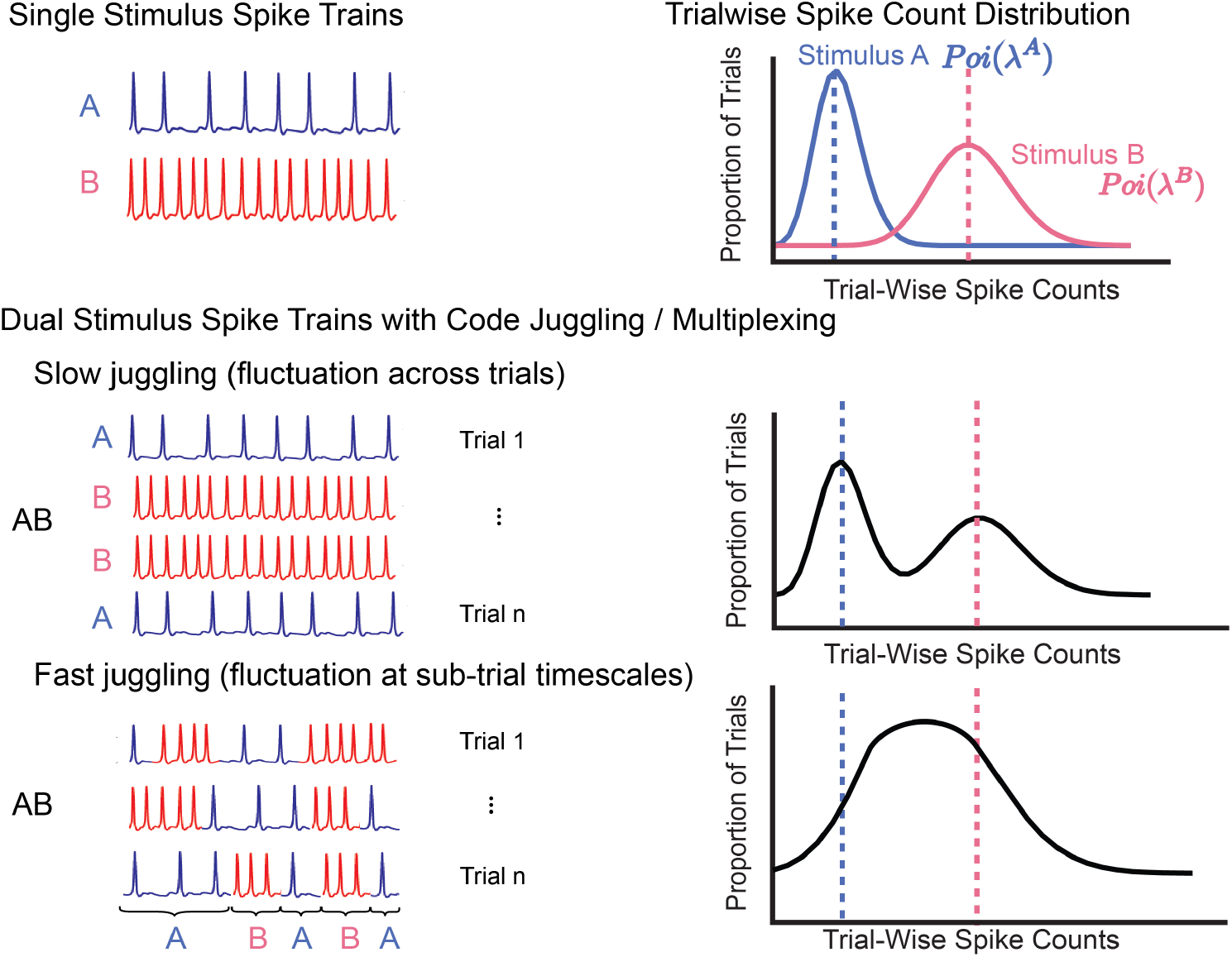
Conceptual framework of code juggling/multiplexing and corresponding distributions of trial aggregated spike count. We consider two scenarios of neural switching patterns when exposed to multiple stimuli. Slow juggling, or fluctuation across trials, indicates switching across trials, leading to a mixture of benchmarked distributions. Fast juggling, or fluctuation at sub-trial timescales depicts switching within trials, where the corresponding distribution concentrates within an interval sandwiched between benchmarked distributions.

Our new method SCAMPI adopts a Bayesian inference framework to test slow and fast juggling against the two controls, all based on trialwise spike count data collected from A, B, and AB exposure trials. The use of a formal Bayesian testing framework, based on Bayes factors, is useful for addressing both whole-trial and sub-trial code juggling and various flavors of alternative behavior. It is also useful to combine single cell evidence into a combined population level inference on the prevalence of code juggling. In our evidence calculation, the Poisson mixture models under both fast juggling and the overreaching alternative are materialized by introduction of continuous mixing distributions which are estimated nonparametrically. The introduction of nonparametric distributions complicates the calculation of Bayes factors. We overcome this issue by adopting the predictive recursion marginal likelihood method of Martin and Tokdar (2011). Our analysis here pertains only to two competing stimuli, but the theory and methods could be generalized in principle to situations with more than two stimuli.

SCAMPI could also be seen as a reinforcement of our previous attempt at evaluating sub-trial switching in which we modeled AB spike trains as a dynamic admixture of point processes (DAPP; Caruso et al. (2018); Glynn et al. (2019)) and carried out statistical estimation by dividing each spike train into smaller bin counts (25-50 ms). While the DAPP analysis can quantify the propensity of a neuron to exhibit whole-trial or sub-trial level switching, or no switching at all, it does not offer a rigorous statistical testing framework to evaluate formal hypotheses about switching. It also faces considerable statistical difficulty in providing precise estimates because of the scarcity of data in short time bins. SCAMPI, which operates with whole-trial level spike count data, can offer much higher precision in evaluating sub-trial switching that is consistent with the fast juggling hypothesis.

## 2 Statistical Analysis Framework

### 2.1 Code Juggling and Alternative Models

Let 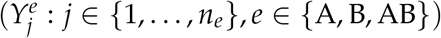 denote trialwise spike count data recorded under three signal settings: exposure to a stimulus A presented in isolation (labeled *e* = A), exposure to a distinct stimulus B presented in isolation (*e* = B), and exposure to stimuli A and B presented together (*e* = AB). The corresponding trialwise spike count distributions are labeled *P*^A^, *P*^B^ and *P*^AB^. Single stimulus spike counts are assumed Poisson distributed:

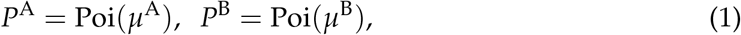

with unknown average firing rates *µ*^A^, *µ*^B^ > 0. The hypothesis of code juggling splits into two sub-type, slow-juggling and fast-juggling:

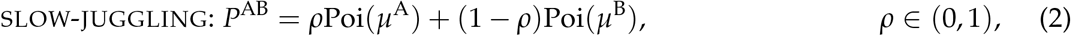

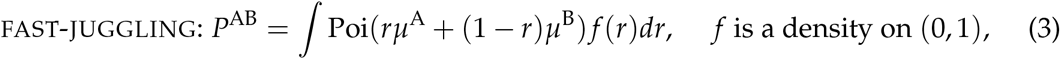

representing, respectively, whole-trial and sub-trial level recombination of *P*^A^ and *P*^B^. slow-juggling captures random, trial-to-trial fluctuation between encoding for A or B at benchmark firing rate of *µ*^A^ or *µ*^B^. Here *ρ* gives the (unknown) relative prevalence of stimulus A within the ensemble. In contrast, fast-juggling captures sub-trial level fluctuations where a random fraction *r* of trial duration is spent encoding for stimulus A; the remaining 1 − *r* fraction is devoted to stimulus B. The fraction *r* quantifies the relative prevalence of stimulus A within the ensemble. It is taken to vary randomly from one trial to the next, according to an unknown density *f* (*r*).

For either code juggling sub-type, *P*^AB^ is relatively overdispersed (variance > mean) compared to *P*^A^ or *P*^B^ (both have mean = variance) and also bracketed between *P*^A^ and *P*^B^ in terms of stochastic ordering^1^. Accordingly, we formalize alternatives to code juggling as distributions that violate either overdispersion or stochastic bracketing, or both.

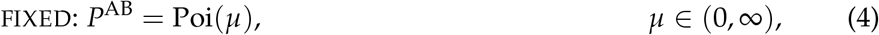

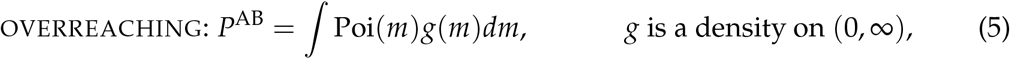

Sub-type fixed represents scenarios where AB is encoded by the neuron as a single fused stimulus of comparable complexity to that of the A or B stimulus. Based on the relative location of *µ* compared to *µ*^A^ and *µ*^B^, we identified four sub-types: (1) fixed-preferred, the case where the neuron always encodes its preferred signal from the ensemble, ignoring the other (*µ* = *µ*^A^ ∨ *µ*^B^); (2) fixed-non-preferred, the case where the neuron always encodes its non-preferred signal (*µ* = *µ*^A^∧ *µ*^B^); (3) fixed-middle, where the neuron encodes a signal that elicits a firing rate between those of benchmarked signals (*µ* ∈ (*µ*^A^∧ *µ*^B^, *µ*^A^ ∨ *µ*^B^)); and (4) fixed-outside, where the neuron encodes a signal that elicits firing rates outside the range of both benchmarked signals (*µ* ∈ (0, *µ*^A^∧ *µ*^B^) ∪ (*µ*^A^ ∨ *µ*^B^, ∞)). The overreaching model is non-benchmarked and overdispersed, representing the remaining scenario where the neuron encodes AB as a more complex or ambiguous signal than either A or B, but the codes may not be related to one another in an information-preservative manner. Instead, AB could simply present as a different type of signal than either component of the ensemble (Festa et al., 2021; Semedo et al., 2019).

Notice that our SCAMPI model encompasses all hypotheses in the model proposed by Caruso et al. (2018), with some nuance (Figure 2). We refer to the model by Caruso et al. (2018) as the original model. Conceptually, slow-juggling would appear to correspond to the mixture category in the original model (Caruso et al., 2018; Mohl et al., 2020; Jun et al., 2022; Schmehl et al., 2024; Groh et al., 2024), but given that assignment to any given category depends on the competition from other categories, the match will be affected by the introduction of the new categories. Specifically, some mixture may now fall into either the fast-juggling or non-benchmarked overreaching categories (Figure 2, Table 1). Conditions that previously fell into the former intermediate category may now be assigned either to fast-juggling or fixed-middle. single is labeled here either fixed-preferred or fixed-non-preferred, depending on which single stimulus “wins”. And the former outside category is now likely to subdivide into fixed-outside and overreaching. In total, the additional categories provide enhanced competition for the old mixture classification, strengthening the robustness of the testing framework in two ways: the odds that overdispersed responses that are not truly related to the individual stimulus distributions are erroneously assigned to the mixture category is reduced, and the possibility of detecting faster fluctuations that would have been overlooked before is improved. This enhancement will be revisited through real analysis, as illustrated in Figure 7.

**Table 1:**
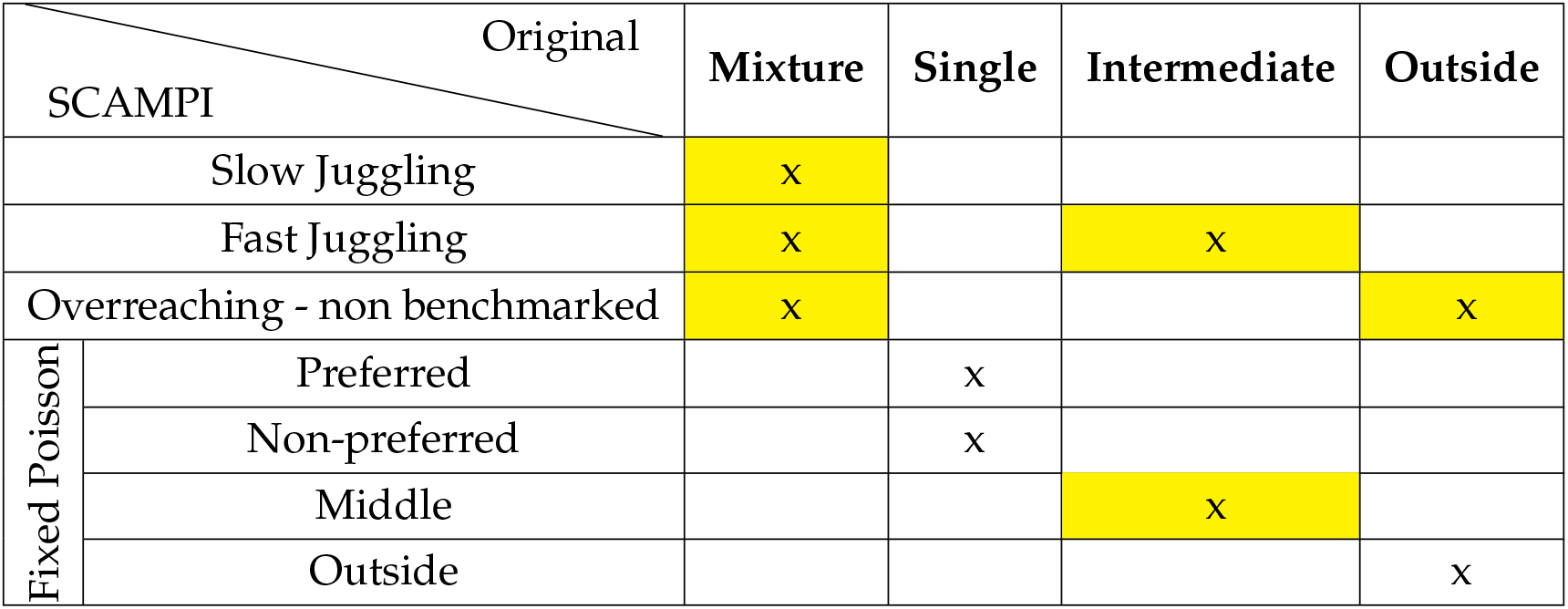
Potential relationship between the whole trial spike count model categories of the original model (columns) and the categories/subcategories of the SCAMPI model (rows). New potential re-assignments that are of particular interest from a neuroscience perspective are highlighted in yellow. For example, the previous mixture category could have included what we now attempt to identify as slow-juggling and fast-juggling, and may also have contained cases better classified as overreaching (overdispersed but unrelated to the benchmarks established from the A-alone and B-alone trials). Similarly, the intermediate category and outside category might encompass multiple distinct categories in the SCAMPI model.

| Original<br>SCAMPI |  | Mixture | Single | Intermediate | Outside |
| --- | --- | --- | --- | --- | --- |
| Slow Juggling |  | x |  |  |  |
| Fast Juggling |  | x |  | x |  |
| Overreaching - non benchmarked |  | x |  |  | x |
| Fixed Poisson | Preferred |  | x |  |  |
|  | Non-preferred |  | x |  |  |
|  | Middle |  |  | x |  |
|  | Outside |  |  |  | x |

**Figure 2:**
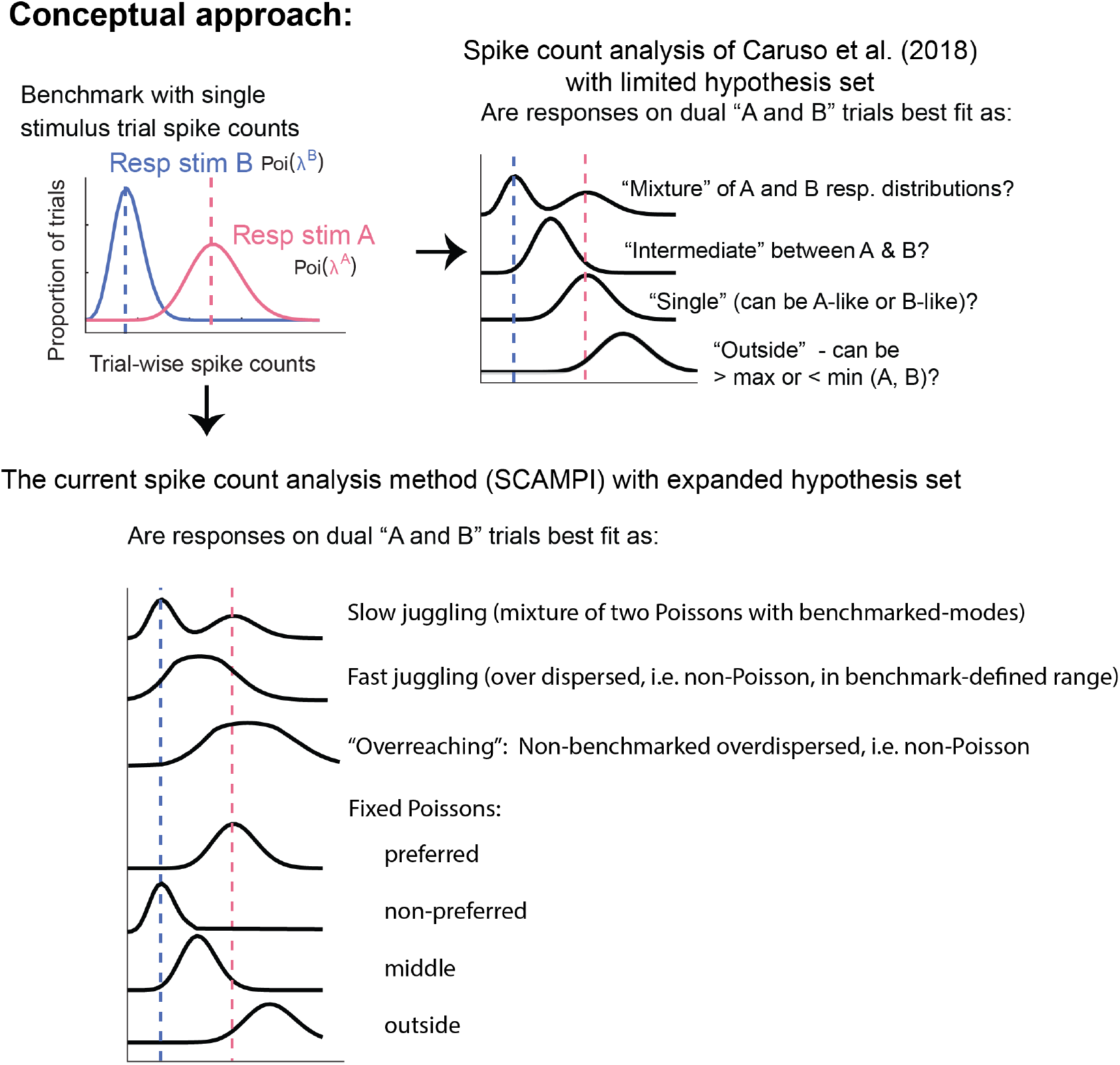
Conceptual approach of the SCAMPI model in comparison to the earlier trialwise spike count model (Caruso et al., 2018; Mohl et al., 2020; Jun et al., 2022; Schmehl et al., 2024; Groh et al., 2024). In both frameworks, we use the number of spikes occurring in response to each stimulus presentation as the metric of activity, obtaining a spike count for each trial. Next, we establish response benchmarks on the basis of the distribution of spike counts observed on A-alone and B-alone trials. In our previous model, we then categorized responses on AB trials into four subgroups, defined in relation to those benchmarks. mixture was defined as the case in which responses appear to be drawn stochastically from either the A-alone or B-alone distributions, varying across trials. The remaining categories were all assumed to be drawn from single Poisson distributions as illustrated. In the new approach, we refine this framework to include additional potential classifications. slow-juggling is nominally identical to our previous mixture category, but the inclusion of a fast-juggling option permits the dissociation of truly across-trial fluctuations from fluctuations that could occur more rapidly – producing an overdispersed distribution that is nevertheless still bounded by the A-alone and B-alone benchmarks. We also include a non-bounded overreaching category, i.e. a response pattern that is broader than a Poisson but does not appear to be constrained by the benchmarks. Finally, there remains a fixed-Poisson category, which can be subdivided into various labels depending on the mean firing rate in relation to the benchmarks. See Table 1 for a summary of the relationship between the old and the new categories.

### 2.2 Nonparametric Mixture Models

All four hypotheses specify the AB spike count distribution as *P*^AB^ = ∫Poi(*µ*^AB^)*dF*(*µ*^AB^) with an unknown rate mixing distribution *F* on (0, ∞), with distinctive requirements on the support of *F* relative to the benchmark rates *µ*^A^ and *µ*^B^. For slow-juggling, *F* is a discrete distribution with support {*µ*^A^, *µ*^B^}, which can be succinctly expressed as 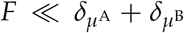, where *δ*_*x*_ denotes the Dirac measure at a point *x* and ≪ refers to absolute continuity of measures. For fast-juggling, 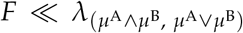 where *λ*_*I*_ denotes the Lebesgue measure restricted to an interval *I* ⊂ ℝ, *u* ∧ *v* = min(*u, v*) and *u* ∨ *v* = max(*u, v*). For overreaching, 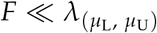, for some prefixed lower and upper bounds 0*≤ µ*_L_ < *µ*_U_ < ∞ on the overall spiking rates. For fixed, we have a degenerate *F* = *δ*_*µ*_ for some unknown *µ* > 0.

A marginal likelihood score for each hypothesis could be obtained by integrating out *F* and *θ* = (*µ*^*A*^, *µ*^B^). Because of the identity *p*(*ϒ*^A^, *ϒ*^B^, *ϒ*^AB^) = *p*(*ϒ*^A^)*p*(*ϒ*^B^) ∫*p*(*ϒ*^AB^|*θ*)*p*(*θ* | *ϒ*^A^, *ϒ*^B^)*dθ*, and the fact that we have the same model for (*ϒ*^A^, *ϒ*^B^) across all scenarios, it suffices to concentrate only on *ϒ*^AB^ as the observed data. The information in (*ϒ*^A^, *ϒ*^B^) can be completely captured through a *second-stage* prior *π*(*θ*) on *θ* obtained as its posterior distribution given (*ϒ*^A^, *ϒ*^B^) under a *first-stage* prior *π*_0_. We choose *π*_0_ to be the Jeffreys’ prior, which is non-informative and invariant under reparametrization. We take 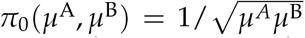, which produces a product gamma distribution in 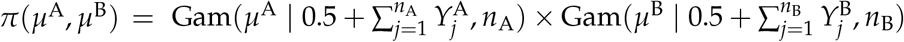.

For the two code juggling hypotheses, i.e., slow-juggling and fast-juggling, we adopt the predictive recursion marginal likelihood (PRML) method of Martin and Tokdar (2011) to integrate out *F* and augment it with a Laplace approximation to integrate out *θ*. These methods are described in the next subsection. In order to apply this framework, one first needs to use a change of variable to express the dependence of *P*^AB^ on *θ* through a kernel *κ*_*θ*_ rather than the support and continuity of *F*. For the slow-juggling case, we can write *P*^AB^(*y*) = ∫ *κ*_*θ*_ (*y*|*u*) *f*(*u*)*dν*(*u*) with *ν* = *δ*_0_ + *δ*_1_, where *κ*_*θ*_ (*y*|*u*) = Poi(*y*| *uµ*^A^ + (1*u*)*µ*^B^), *u* ∈ {0, 1}. The same kernel works for the fast-juggling case, but now with *u*∈ (0, − 1) and *ν* = *λ*_(0,1)_.

In the case of overreaching, the marginal likelihood involves an integration of *F* alone. No integration over *θ* is needed since the support of *F* is free of *θ*. We approximate the integral over *F* via PRML with the kernel *κ*(*y* | *u*) = Poi(*y* | *uµ*_L_ + (1 − *u*)*µ*_U_), each with *u* ∈ (0, 1) and *ν* = *λ*_(0,1)_. An initial guess *f*_0_ of *f* is needed to carry out PRML in each of these three cases. We take *f*_0_ to be the appropriate uniform density with respect to the corresponding dominating measure *ν*; see Table 2.

**Table 2:** The SCAMPI model settings under different hypotheses on *P*^AB^ in a reparametrized form. Notice that the fixed hypothesis is not listed here since we can calculate marginal likelihood by integrating out *µ*.

| Model | $\kappa_\theta(y \mid u)$ | Support | $f_0(u)$ | $\nu$ |
| --- | --- | --- | --- | --- |
| SLOW-JUGGLING | $\text{Poi}(y \mid u\mu^A + (1-u)\mu^B)$ | $u \in \{0, 1\}$ | 0.5 | $\delta_0 + \delta_1$ |
| FAST-JUGGLING | $\text{Poi}(y \mid u\mu^A + (1-u)\mu^B)$ | $u \in (0, 1)$ | 1 | $\lambda_{(0,1)}$ |
| OVERREACHING | $\text{Poi}(y \mid u\mu_L + (1-u)(\mu_U))$ | $u \in (0, 1)$ | 1 | $\lambda_{(0,1)}$ |

Finally, the marginal likelihood score for fixed could be calculated via direct integration over the scalar parameter *µ*. Again, no integration over *θ* is necessary. The integration over *µ* must be carried out under a suitable prior specification. A reasonable default choice is the Jeffreys’ prior *π*(*µ*) = *Cµ*^−1/2^, defined up to an arbitrary constant *C* > 0. This presents a technical difficulty for marginal likelihood calculation, because the integral is also defined up to the arbitrary multiplicative constant *C*. A potential remedy is to borrow from the intrinsic Bayes factor proposal of Berger and Pericchi (1996) where marginal likelihoods are replaced with intrinsic marginal likelihoods. Specifically, a subset of the data is sacrificed to update (*train*) the improper prior into a proper probability distribution, which is then used to calculate the marginal likelihood of the remaining data. Berger and Pericchi (1996) advocate carrying out the intrinsic adjustment to all hypotheses under consideration, and using the same training subset across models. We do not pursue this approach because the intrinsic adjustment to the other three models substantially adds to the computational cost. We experimented with a cost-effective modification of this proposal where the intrinsic adjustment was applied only to fixed, but the resulting tests were found to be heavily biased in favor of fixed in synthetic data experiments (Supplementary Figure 11). A second remedy is to truncate Jeffreys’ prior to the compact interval [*µ*_L_, *µ*_U_] used for overreaching. We do not pursue this either because our simulation studies showed that the resulting tests are heavily biased against fixed (Supplementary Figure 12).

Instead, we focus on fixing *C* to ensure that the probability of incorrectly rejecting fixed is controlled at a given threshold. In other words, if one fixated on our test as an assessment of fixed versus the other hypotheses, what *C* do we need to ensure a given *α* level of significance? We established through a numerical study (Supplementary Figure 10) that the choice of *C* = 1 ensures an *α* = 5%. All our numerical analyses are carried out with the choice of *C* = 1.

### 2.3 Predictive Recursion Marginal Likelihood and Bayes Factor

Suppose data *ϒ*_1_, …, *ϒ*_*n*_ are modeled as independent realizations from the mixture distribution

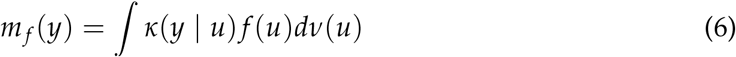

where (*y, u*) ↦ *κ*(*y*|*u*) is a known kernel on 𝒴 × 𝒰 and *f* is an unknown mixing density in 𝔽: the set of probability densities with respect to a *σ*-finite Borel measure *ν* on 𝒰. The predictive recursion algorithm (Newton et al., 1998) estimates *f* by starting with an initial guess *f*_0_ ∈ 𝔽 and then making a single pass through the observations to recursively update to a final estimate *f*_*n*_. At stage *i* ∈ {1, …, *n*} of the recursion, only observation *ϒ*_*i*_ is used to update the current estimate *f*_*i*−1_ to a new estimate *f*_*i*_ according to,

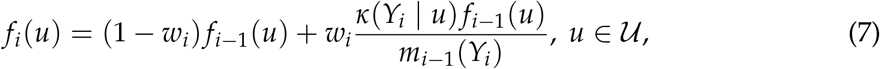

where *w*_*i*_ ∈ (0, 1) are prefixed recursion weights and

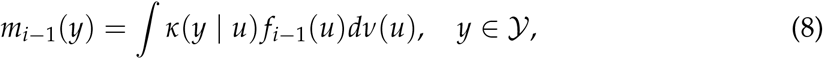

is the estimated predictive density of *ϒ*_*i*_ given *ϒ*_1_, …, *ϒ*_*i*−1_. Under certain regularity conditions on the kernel *κ*(*y* | *u*), both *f*_*n*_ and *m*_*n*_ asymptotically converge to the true densities *f* = *f*^⋆^ and *m* = *m*^⋆^ := *m*_*f* ⋆_, respectively, provided 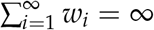 and 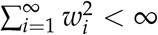 (Tokdar et al., 2009). The regularity conditions remain valid in our case, wherein we utilize Poisson kernels with mean parameters restricted within a compact space.

When the kernel *κ* = *κ*_*θ*_ involves an unknown parameter *θ* ∈ Θ, Martin and Tokdar (2011) recommend using the PRML score

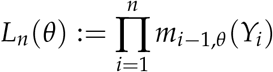

to evaluate a marginal likelihood score of *θ*, which is easily computed as a by-product of running the predictive recursion algorithm with *θ* fixed at the given value. This recommendation is justified on three accounts. First, since *m*_*i*−1,*θ*_ gives an estimate of the predictive density of *ϒ*_*i*_ given *ϒ*_1_, …, *ϒ*_*i*−1_ and *θ*, the proposed likelihood score gives a corresponding estimate of the true marginal density evaluation 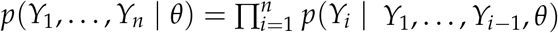. Second, under appropriate regularity conditions, the PRML score enjoys the asymptotic consistency property that *n*−1 log{*L*_*n*_(*θ*)/*L*_*n*_(*θ*^⋆^)} ↦ − inf _*f* ∈ 𝔽_ *d*_KL_(*m*^⋆^, *m*_*f*, *θ*_). Third, for the special choice of *w*_*i*_ = (1 + *a*)^−1^, for some *a* > 0, predictive recursion could be seen as an expectation filtration approximation to a fully Bayesian estimation of *f* under a Dirichlet process prior with precision *a* and base density *f*_0_, and hence, *L*_*n*_(*θ*) could be seen as the corresponding filtration approximation to the fully Bayesian marginal likelihood score of *θ*. Martin and Tokdar (2011) provide extensive numerical evidence of the latter correspondence.

Now consider a finite set of competing mixture models 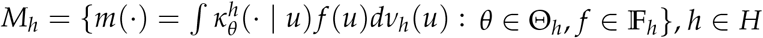 We propose to evaluate a Bayes factor *B*_*i,j*_ between models *M*_*i*_ and M_j_ as B_12_ = I(M_i_)/I(M_j_) where

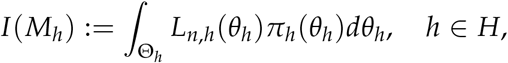

gives the integrated PRML score under an appropriate prior *π*_*h*_ on Θ_*h*_. Again, our interpretation of this ratio as a Bayes factor relies on interpreting each *L*_*n,h*_(*θ*) as an approximation to the marginal likelihood score under the corresponding Dirichlet process mixture idealization of *M*_*h*_. Extending this analogy we also propose computing a posterior probability for model *M*_*h*_ as:

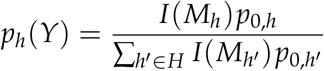

where *p*_0,*h*_, *h* ∈ *H* are prior weights assigned to the competing hypotheses. The hypothesis with the highest posterior probability is the winning model, where the posterior probability measures the confidence of making a right choice from these four hypotheses suggested by the data. We consider 0.25− 0.5, 0.5− 0.75, 0.75− 1.0 as the weak, moderate and strong confidence on the model choice respectively.

Assuming that the kernel 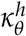 is smooth in *θ* allows us to approximate *I*(*M*_*h*_) by a Laplace type approximation: 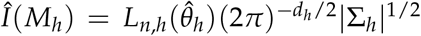, where 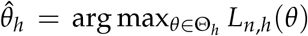, and 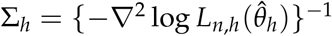. The necessary optimization could be carried out by using the gradient based extension of PRML in Martin and Tokdar (2011) in which 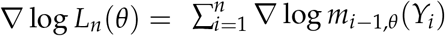 is calculated recursively during the same single pass through the data used for the original predictive recursion and PRML calculation. In our applications, we use the gradient-based optimzation algorithm due to Broyden-Fletcher-Goldfarb-Shanno (BFGS) to compute 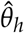. Then, a numerical evaluation is done of Σ_*h*_.

It is worth noting that the PRML score *L*_*n*_(*θ*) depends on the order in which the observations enter the single-pass recursive algorithm. Such an order dependence is problematic from both theoretical and practical perspectives when data are believed to be exchangeable. We could mitigate this problem by considering a permutation version of the PRML score that averages over the *n*! many scores obtained by running the single pass algorithm on every possible permutation of the data. In practice, however, it suffices to consider a Monte Carlo approximation, where averaging is done on a subset of randomly drawn permutations.

## 3 Performance Assessment

### 3.1 Experimental Design

To assess statistical accuracy of the SCAMPI model, we conducted experiments using synthetic data and focused on two key aspects: detecting code juggling, and correctly identifying *P*^AB^ as one of fixed, slow-juggling, fast-juggling, and overreaching. We generated data from four scenarios: F, SJ, FJ, and O, which correspond to four hypotheses: fixed, slow-juggling, fast-juggling, overreaching, respectively. These four scenarios have a tuning parameter controlling the complexity of the classification task. For each scenario, 100 experiment sets were created, each set comprising of synthetic A, B and AB trial data with *n*_A_ = *n*_B_ = *n*_AB_ ∈ {20, 30, 50} Trial sizes and average A and B firing rates were set to broadly match similar statistics obtained from IC data described in Section 4. For every experiment set, A and B spike counts were generated, respectively, from *P*^A^ = Poi(50) and *P*^B^ = Poi(80).

The cases differed from one another in the choice of *P*^AB^ and the complexity parameter. The exact form of data generating models *P*^AB^ are:

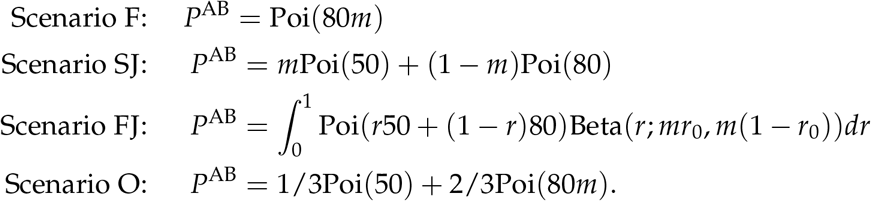

A visual summary is given in Figure 3. For scenario F, complexity is coded as a multiplier *m* to the mean. The multiplier *m* is varied to encompass all sub-types of the fixed hypothesis: fixed-preferred, fixed-non-preferred, fixed-middle, fixed-outside. With

**Figure 3:**
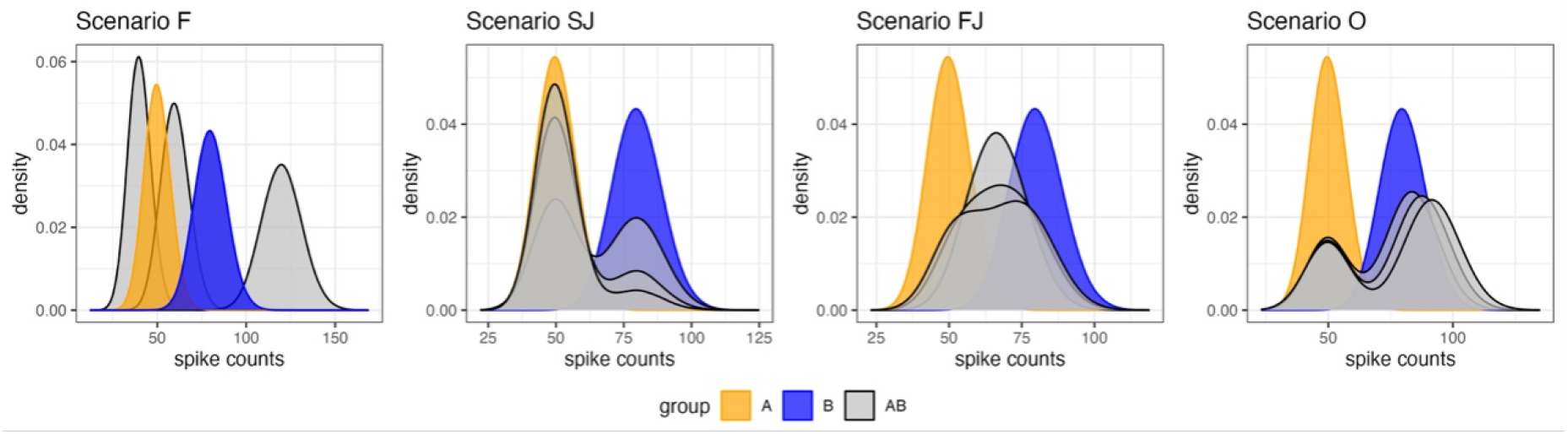
Choice of (*P*^A^, *P*^B^, *P*^AB^) for all groups in the experimental design. Grey graphs show *P*^AB^.

*m* = 5/8, *P*^AB^ equals *P*^A^. Similarly, with *m* = 1, *P*^AB^ = *P*^B^. Notice that *P*^AB^ = *P*^A^ and *P*^AB^ = *P*^B^ are boundary cases for each of the other three hypotheses. For example, setting *ρ* = 1 in the mixture formulation (2) produces *P*^AB^ = *P*^A^. Therefore *m* = 5/8 or *m* = 1 corresponds to ambiguity between fixed and other hypotheses, making the classification task more complex for *m* values near these two points. In addition, if *m* falls between these two points, where *m* ∈ (5/8, 1), the corresponding firing rate *µ*^AB^ will be bounded by the firing rates of constituent stimuli *µ*^A^ and *µ*^*B*^. As a result, the corresponding probability distribution *P*^AB^ becomes a borderline case of fast-juggling, which also poses a great challenge for classification.

For scenario SJ, complexity increases as the skewness parameter *m* gets close to either 0 or 1, making *P*^AB^ close to the boundary cases, respectively, *P*^A^ or *P*^B^.

For scenario FJ, where a beta distribution with mean *r*_0_ is used as the mixing density, complexity is controlled by varying the precision parameter *m*, which in turn controls the fano factor of the resulting *P*^AB^. We take *r*_0_ = 0.56 and let *m* vary from zero to infinity so that the fano factor goes from a minimum possible value of 1 to a maximum possible value of 4.5. When the fano factor is 1, *P*^AB^ = Poi(63.2) and hence can also be explained as fixed. When the fano factor is at the maximum, *P*^AB^ = 0.56Poi(50) + 0.44Poi(80), causing confusion with slow-juggling. So the complexity of the task is high at both extremes of the fano factor values. See Supplementary Materials for more explanation of the choice of *r*_0_ and the range of fano factor values one can obtain with a beta mixing density.

For scenario O, we consider bimodal mixtures of Poi(*µ*^A^) with another Poi(*mµ*^B^), with the multiplier *m* controlling complexity. As *m* moves closer to 1, *P*^AB^ reduces to a mixture of Poi(*µ*^A^) and Poi(*µ*^B^) and hence can be explained by the slow-juggling hypothesis, making a correct classification of overreaching difficult.

For each scenario, PRML based posterior probabilities of the four competing hypotheses were calculated. For each set, we recorded whether the correct label received the maximum posterior probability, i.e., whether a correct identification was made. Additionally, we recorded whether the test correctly detected code-juggling. It’s important to note the distinction between correct classification and correct detection of code-juggling. If the truth were slow-juggling, being classified as fast-juggling will still be counted as a correct detection of code-juggling even though it is a misclassification. Similarly, when a true fast-juggling case is misclassified as slow-juggling, it would still count toward a correct detection of code-juggling. In other words, when the focus is detection of code-juggling, a correct identification of the exact sub-type is viewed as of secondary importance.

### 3.2 Results

Accurate detection of code-juggling is the primary focus for our applications. We consider two key metrics: sensitivity, which is the proportion of code-juggling cases that are correctly identified as code-juggling, and specificity, which is the proportion of non-code-juggling cases that are correctly identified as not being code-juggling. Figure 4 demonstrates that our PRML test performs well in terms of sensitivity and specificity. For data generated from scenario F, specificity is above 0.9, even for cases where the multiplier *m* is close to either 1 or 5/8, or falls between these two points. For data generated from scenario O, even with a small sample size of 20, specificity can reach 0.75 if the multiplier *m* is greater or equal to 1.15, where one of two modes is reasonably away from *µ*^B^. As the sample size increases, specificity improves significantly when the multiplier *m* ranges between 1.1 to 1.15. The sample size has less impact on specificity as the multiplier *m* approaches 1. For data generated from scenario SJ, sensitivity is above 0.8 if the skewness parameter is less than 0.85. Increasing the sample size to 50 improves sensitivity to be above 0.75 even for the complicated cases, where the skewness parameter is equal to 0.95. When the true generating process is scenario FJ, sensitivity is above 0.85, if the fano factor is greater than or equal to 2.5. However, as the fano factor approaches 1, more fast-juggling triplets are mis-classified as fixed, lowering sensitivity. Increasing the sample size to 50 mitigates this issue to some extent, ensuring sensitivity is at least 0.625 even for complex cases, where the fano factor is equal to 1.5. It is worth noting that as the fano factor approaches 4.5, more fast-juggling triplets are mis-classified as slow-juggling, which is still a sub-type of code-juggling, and thus may not be of much practical concern.

**Figure 4:**
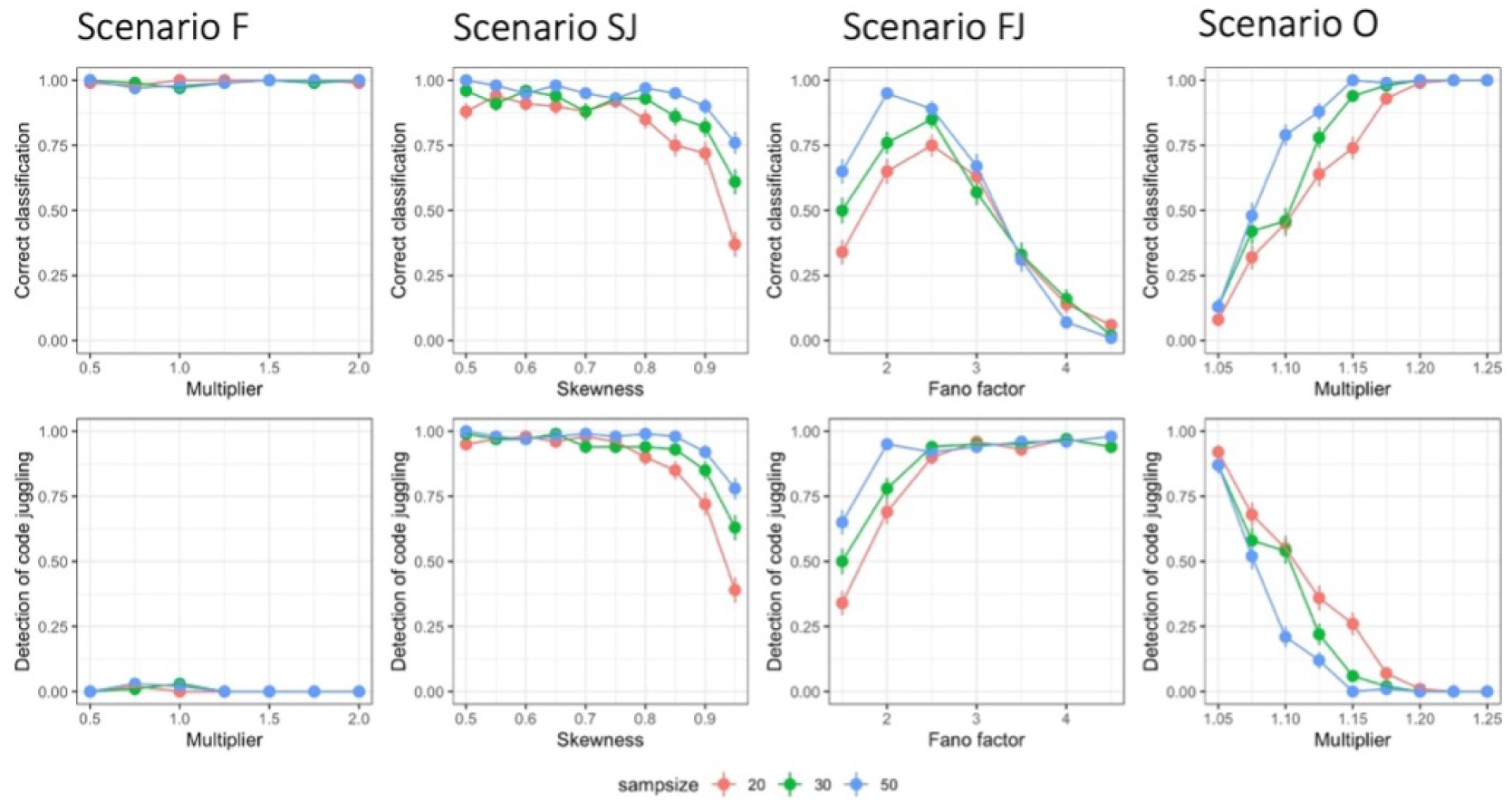
Classification accuracy and code juggling detection accuracy versus complexity parameters across fixed, slow-juggling, fast-juggling, and overreaching experimental cases (columns). Sample size 20, 30, 50 are color coded as red, green and blue.

The most challenging cases are found at the limits of the model spaces, where they lie at the boundary of four hypotheses. These cases can be categorized into three main types. Firstly, sensitivity is low for scenario SJ with a high skewness or for scenario FJ with a low fano factor, where it tends to be mis-classified as fixed. Secondly, specificity is low for scenario O with a low multiplier, where it tends to be mis-classified as slow-juggling. Thirdly, there is ambiguity between sub-types of code-juggling for scenario FJ with a high fano factor, since it tends to be classified as slow-juggling. As explained earlier, the model spaces underlying the four hypotheses are nested in the order: fixed *⊂* slow-juggling *⊂* fast-juggling *⊂* overreaching. For aforementioned complicated situations arising at the boundaries of the model spaces, a simpler model is always favored by the PRML test due to Occam’s razor property of Bayes factor. Increasing the sample size can greatly improve the sensitivity for the first type of challenge, but has very limited impact on specificity for the second type of challenges, especially when cases are near the boundaries. Additionally, increasing the sample size does not help in identifying sub-types of code juggling, the third type of challenge. As previously noted, distinguishing between two sub-types of code juggling is not a primary concern in practical applications.

## 4 Applications

Caruso et al. (2018) first reported statistical evidence espousing the viewpoint that an individual neuron may display fluctuating activities when preserving dual stimuli in visual and auditory areas, such as the inferior colliculus (IC) area and the middle fundus (MF) face patch systems in the inferotemporal (IT) cortex. Subsequently, such fluctuating patterns were observed in other visual areas, including the primary visual cortex (V1), visual area V4, the middle temporal visual area (MT), and the anterior lateral (AL) face patch systems in the IT cortex (Jun et al., 2022; Schmehl et al., 2024). Moreover, fluctuations in V1 and MT have been demonstrated to relate to object identification (Jun et al., 2022; Schmehl et al., 2024). However, as we mentioned in Section 1 earlier, our original analysis method could not distinguish fast vs. slow juggling, and fluctuations that were not related to the single stimulus benchmarks could have been labeled as juggling when such a categorization may not be appropriate. Accordingly, we reanalyzed the IC, V1, and IT datasets from the earlier studies, to (1) confirm the presence of code juggling; (2) determine what proportions may be fast vs. slow, (3) validate our earlier findings that code juggling is more prevalent for combinations of faces than for face-object pairs in the face patch system (Schmehl et al., 2024); and is associated with object identification (Jun et al., 2022) for individual neurons in the V1 area.

### 4.1 Datasets

#### IC dataset

Caruso et al. (2018) reported a study where spiking activity of individual IC neurons were recorded while monkeys made saccades towards one or two sound locations (Figure 5(A)). In dual-sound trials, the two sounds were presented in different hemifields, and consisted of two distinct frequencies (differing in frequency by at least 22 %). Every dual-sound condition was matched to two single-sound conditions corresponding to each of the two constituent location-frequency pairs. Stimulus conditions were shuffled within and across experiment sets. Here, the spike counting window was between 0 ms and 600 ms (the original study used some longer time windows).

**Figure 5:**
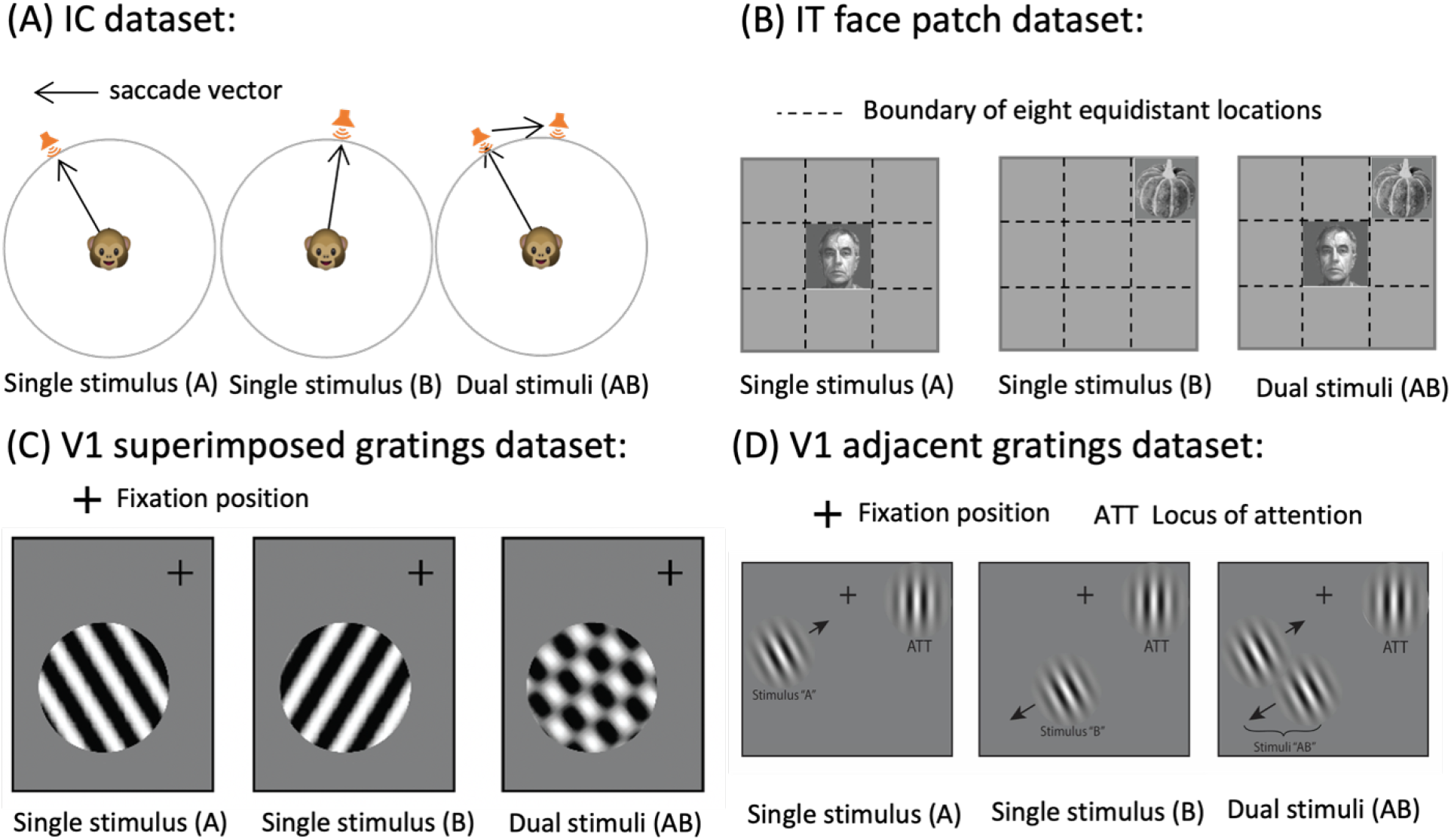
Stimulus conditions for the datasets re-analyzed in the current study. (A) For the IC dataset, bandpass sounds of different center frequencies were presented individually or simultaneously at different spatial locations, and monkeys made saccades to any and all sounds presented. No constraints were imposed on which sound the monkey looked at first. See Caruso et al. (2018) for details. (B) In the face-patch datasets, stimuli consisted of either faces or non-face objects presented at different locations. Monkeys performed a fixation task. See Ebihara (2015); Caruso et al. (2018); Schmehl et al. (2024) for details. (C) In the V1 superimposed gratings dataset, one or two gratings were presented at the same location while monkeys fixated. See Jun et al. (2022) and Ruff and Cohen (2016) for details. (D) In the V1 adjacent gratings dataset, the A and B conditions consisted of single gratings presented in the contralateral hemifield while monkeys attended to another grating presented in the ipsilateral hemifield. The AB condition involved both of those A and B gratings presented side-by-side while monkeys continued to attend to a third grating in the ipsilateral hemifield. See Jun et al. (2022); Ruff and Cohen (2016) for details.

#### IT face patch dataset

In a study by Ebihara (2015), spike activities of individual cells in the MF and AL face patch areas were recorded when monkeys performed a fixation task (Figure 5(B)). The stimuli presented were classified into two categories: a preferred face which elicited a high firing rate, and a non-preferred stimulus (either a face or an object) which elicited little to no neural activity. During dual stimuli trials, a preferred face was always presented at the center of the receptive field, with the non-preferred stimulus presented adjacent to it simultaneously at varying locations. In corresponding single stimulus trials, one of the constituent stimuli was presented alone. Although different faces and objects were considered as different experimental conditions, the heterogeneity of the locations of non-preferred stimuli was ignored, following the analysis by Caruso et al. (2018). The spike counting window was set between 50 ms and 250 ms (same as before).

#### V1 superimposed gratings dataset

Ruff et al. (2016) utilized a multi-electrode array to record spike activities of multiple cells in the V1 area while monkeys passively viewed drifting gratings (Figure 5(C)). In dual stimuli trials, superimposed drifting gratings with orthogonal orientations and moderate contrast were presented, producing a “plaid” appearance. In corresponding single stimulus trials, the component gratings were presented individually. Distinct orientations of gratings defined different experimental conditions. Following the analysis of Jun et al. (2022), the spike counting window was between 30 ms and 230 ms.

#### V1 adjacent gratings dataset

Similar to the superimposed grating dataset, Ruff and Cohen (2016) also used multi-electrode recordings in V1 area, but their stimuli consisted of smaller drifting Gabor patches (Figure 5(D)). For the data included in the present study, either one grating or two gratings were presented in the contralateral hemifield, covering the set of receptive fields of the V1 neurons, while an additional grating was always presented in the ipsilateral hemifield. Monkeys performed a motion direction change detection task involving that ipsilateral stimulus. We treated trials with one contralateral grating as single stimulus trials and two contralateral gratings as dual stimulus trials. Following Jun et al. (2022), the spike counting window was taken to be 30 ms through 230 ms.

### 4.2 Preprocessing

We reanalyzed the same data with the SCAMPI model with one additional difference in the preprocessing step. Caruso et al. (2018) tested the Poisson distribution assumption on A and B spike counts by using a chi-square goodness of fit test which calculated expected and observed counts by binning the data histogram. This test has the ability to detect both over- and under-dispersion. For the purposes of our analysis, overdispersion of *P*^A^ and *P*^B^ is a bigger issue than underdispersion. If trial-to-trial variability was caused by factors extrinsic to the chosen stimuli, one would expect overdispersed count distributions under all three conditions.

We replaced the chi-square goodness of fit test with thresholding fano factor (variance-to-mean ratio) greater or equal to 3 to remove the triplets that had heavily over-dispersed distributed *P*^A^ and *P*^B^ (Schmehl et al., 2024). By filtering out triplets where either *P*^A^ or *P*^B^ is overdispersed, we secured a more conservative position in interpreting classification analysis results of the remaining triplets purely through the lens of stimulus-related code juggling. Thresholding fano factor at 3 retains slightly more triplets compared to the screening based on Poisson variance test with p-value greater than 0.001.

Keeping with other preprocessing steps of previous studies (Caruso et al., 2018; Jun et al., 2022; Schmehl et al., 2024), we only included triplets having at least 5 trials for each condition (A, B, AB), and with well-separated^2^ single-stimulus distributions *P*^A^ and *P*^B^. Summary statistics for each dataset are shown in Supplementary Figure 8.

### 4.3 Results and Interpretation

We applied the SCAMPI model on the preprocessed triplets to first classify the triplets into four categories, namely fixed, fast-juggling, slow-juggling, and overreaching. To permit comparisons with earlier findings (e.g. Caruso et al. (2018)), we also further sub-classified the fixed triplets (*P*^AB^ = Poi(*µ*), *µ* ∈ (0, ∞)) into the “preferred”, “nonpreferred”, “middle” and “outside” categories (Figure 2, Table 1). Triplets were classified according to the highest posterior probability, which also quantifies the level of confidence in the classification. A full report of the number of triplets categorized in each category by dataset and confidence level is provided in Supplementary Figure 9 and Table 3. To facilitate comparison across datasets, we calculated the percentage of occurrences for each “winning” category within a given dataset, subsequently comparing these percentages across datasets within the same “winning” category.

**Table 3:** Total count for different classification and datasets by the SCAMPI model.

|  | Classification by SCAMPI Model |  |  |  |  |  |  | Total |
| --- | --- | --- | --- | --- | --- | --- | --- | --- |
|  | FastJug | SlowJug | OvReach | FxdPrf | FxdNon | FxdMid | FxdOut |  |
| IC | 55 | 25 | 55 | 79 | 45 | 155 | 28 | 442 |
| IT-AL | 26 | 0 | 48 | 65 | 2 | 18 | 0 | 159 |
| IT-MF | 24 | 0 | 21 | 79 | 0 | 52 | 1 | 177 |
| V1-imp | 9 | 3 | 39 | 869 | 4 | 48 | 736 | 1708 |
| V1-adj | 75 | 57 | 171 | 1372 | 4 | 216 | 40 | 1935 |

Results for category assignments are illustrated in Figure 6, focusing first on the two “code juggling” categories, fast and slow (Figure 6(A)). These results reaffirm our earlier findings that code juggling is a general coding scheme observed in the IC area, IT area, and V1 area (Figure 6(A)). In the IC, fast-juggling and slow-juggling together accounted for a non-negligible proportion of the classifiable – about 18.2% of such triplets. In contrast with our earlier evaluation of this dataset, the new analysis method suggests that fastjuggling is more common than slow-juggling (blue bar, ∼12.5%, vs orange bar, ∼5.7%).

**Figure 6:**
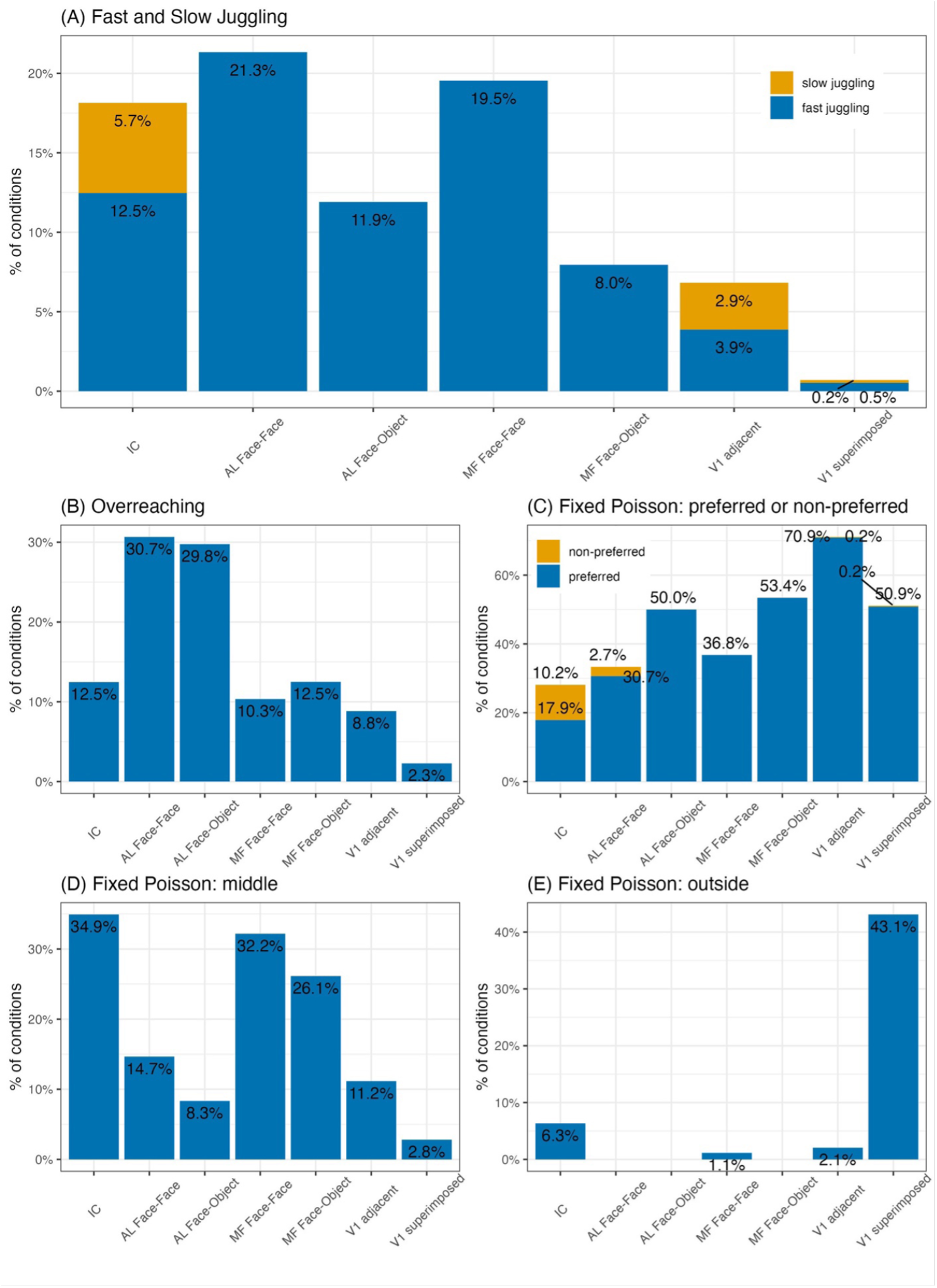
Results of the SCAMPI model for the IC, IT face patch, and V1 datasets. Percentages shown reflect the percentage of conditions, out of the total number of conditions. (A) Prevalence of fast-juggling and slow-juggling across datasets. fast-juggling is seen in non-negligible amounts in all datasets except the V1 “superimposed grating” dataset. It is generally more common than slow-juggling. Juggling is more common for face-face than for face-object datasets in the facepatch system. (B) Prevalence of overreaching. This was most common in AL. (C) Prevalence of fixed poisson (preferred and non-preferred). Conditions were far more likely to be classified as fixed-poisson-preferred than fixed-poisson-non-preferred. (D) Prevalence of Fixed-Poisson-middle. Such a response pattern may indicate that neurons are averaging their inputs. (E) Prevalence of fixed-poisson-outside. This pattern was chiefly observed in the V1-superimposed grating dataset.

In IT cortex, fast-juggling was observed – at roughly equivalent levels – in both AL and MF when two faces were presented (21.3% and 19.5% respectively), and in both areas the prevalence of fast-juggling was higher for two-faces than for face-object combinations (11.9% and MF: 8.0% respectively). This supports the suggestion we previously made in Schmehl et al. (2024) that selectivity for the type of stimulus involved may contribute to the prevalence of juggling patterns. In the case of the face patch system, non-face objects are not thought to evoke strong responses; rather, a distinct population of neurons outside these areas are likely to be encoding these stimuli and “juggling” may therefore not be needed within the face patches when only one of the two stimuli is a face.

In V1, 6.8% of the triplets were categorized as “juggling” when adjacent stimuli were presented. Of these, fast-juggling (3.9%) was slightly more common than slow-juggling (2.9%). Although “juggling” was seen at lower levels than in the IC and IT cortex regions, the relative rarity of such switching potentially relates to the much smaller size of the receptive fields in this structure. Small receptive fields mean different stimuli can potentially be encoded by separate subpopulations of neurons, reducing the need for “juggling” within neurons.

In contrast, “juggling” was much less prevalent in the V1 superimposed gratings experiment, where Gabor patches were presented as a fused object (a plaid, see Figure 5(C)): only 0.7% of triplets were labeled as code juggling. This confirms our previous observation that “juggling” appears to be connected to perceptual object formation (Jun et al., 2022; Schmehl et al., 2024; Groh et al., 2024).

Across all the datasets and conditions, fast-juggling was more prevalent than slowjuggling. This supports our earlier suspicion that the previous mixture category, while nominally identifying switching activity across trials, included many cases of activity where some “switches” actually happened within trials (We will discuss in more detail later, as shown in Figure 7 for an example). Our current results suggest that such within trial fluctuation is indeed common.

**Figure 7:**
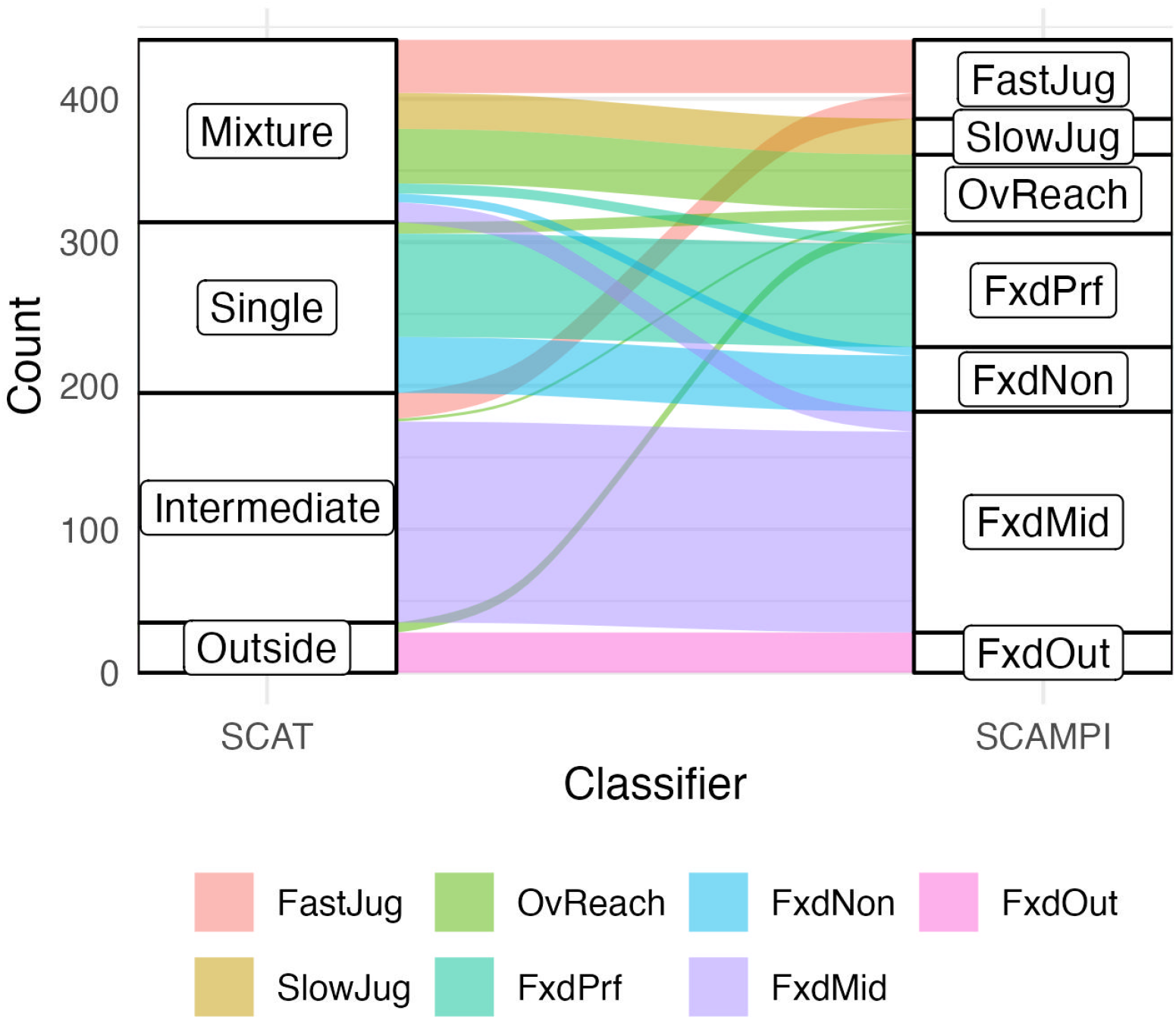
Results from both the original model and the SCAMPI model for the IC dataset. Each stratum presents different categories under distinct frameworks, with the height indicating the frequency of triplets classified into each category. Each stream represents a triplet, flowing from the classification of the original model to that of the SCAMPI model. Classifications under the SCAMPI model was largely consistent with the original model. A significant proportion mixture triplets was classified as overreaching, suggesting that the detected fluctuations were unrelated to encoding benchmarked stimuli. Moreover, more mixture triplets were classified as fast-juggling than slow-juggling, indicating a higher prevalence of fast-juggling in the IC dataset. Additionally, a non-negligible proportion of intermediate triplets was classified as fast-juggling, suggesting that triplets previously assumed to average between two benchmark stimuli may actually switch between them at a rapid rate. The SCAMPI model unravels previous ambiguous cases to a more refined scale.

The winning percentages of the other model categories varied across studies. overreaching was relatively common among the studies that showed signs of juggling, but rare in the V1 superimposed dataset (2.3%) (Figure 6(B)). Recall that overreaching is indicative of fluctuating activity under dual-stimuli exposure, just that the patterns of these fluctuation are not demarcated by the A and B benchmarks. It is entirely possible that some overreaching patterns represent a form of code juggling but one in which the spike count benchmarks observed on single-stimulus trials do not yield good predictions of those found on dual-stimulus trials. One possibility is that an attentional process comes into play on dual stimulus trials, adjusting the signal strength for each stimulus and changing the expected benchmark values. Another possibility is that a more subtle operation involving spike timing is at play, which analyses at the spike count level overlook; a finer timescale analysis of the underlying spike trains would be necessary. Such directions remain interesting avenues for future explorations.

Across all studies, a common occurrence was neurons responding to just one stimulus or another (fixed-preferred and fixed-non-preferred); see Figure 6(C). Such responses are information-preserving, since only one stimulus is encoded in a given neuron’s activity pattern. Whereas previously we did not distinguish between “winner-take-all”, i.e., responding at a level corresponding to the preferred stimulus, and “loser-take-all”, i.e., responding at a level corresponding to the non-preferred stimulus, we can now see that the majority of response patterns across datasets involve encoding the preferred stimulus (Figure 6(C) blue bars larger than orange bars).

Membership in the categories fixed-middle and fixed-outside varied considerably across datasets (Figure 6(D) vs Figure 6(E)). fixed-middle suggests that an averaging operation is at play, i.e., neurons may be responding at a value corresponding to the average of their A and B-like responses. This was more common in the IC and the MF face patch than in the other datasets. However, this pattern can also occur if there is switching between signaling of A and B that is both fast and regular. That is, if neurons switch rapidly between encoding A and encoding B within the spike-counting window, and if the amount of time spent encoding A and encoding B is consistent across trials, then the resulting spike count is likely to be well fit by a single Poisson with a mean rate that is in the middle between the A and B rates. Indeed, the DAPP model of Caruso et al. (2018) and Glynn et al. (2019) identified patterns of rapid fluctuation for these two datasets.

fixed-outside was the predominant outcome in the V1 superimposed datasets, accounting for 43.1% of the classifiable conditions. This observation suggests that neurons may treat the fused object as a new object unrelated to its constituent stimuli in multiple trials. In contrast, in the V1 adjacent gratings dataset, where Gabor patches were presented adjacent to each other (Figure 5(D)), only 2.1% of triplets are labeled as fixed-outside, while 6.8% of triplets are labeled as code juggling, which is significantly different from the rate under superimposed conditions.

### 4.4 Comparison of classifications between the original Caruso et al. (2018) and SCAMPI models

Here we turn to a detailed comparison of the SCAMPI findings with those of the earlier analyses which were primarily focused on the slow-juggling hypothesis. We use the IC study as a representative example (refer to the alluvial plot Figure 7) to showcase the mapping between the two frameworks. Results for the remaining studies are provided in the Supplementary Figure 14 and Figure 13.

In Figure 7, the left and right columns depict classifications under the original model and SCAMPI model, respectively. Each stratum displays different categories under different frameworks, with the height representing the frequency of triplets classified into a particular category. Each stream represents a triplet, colored according to its classification under the SCAMPI model. A stream flowing from mixture to fast-juggling indicates that the triplet was classified as mixture in the original model but as fast-juggling in SCAMPI model.

We note that ambiguity primarily occurred within mixture and occasionally in intermediate categories, consistent with Table 1. As intended, the mixture category of the original model now subdivides into several classifications in the SCAMPI model. About half of previously labelled mixture triplets retain a multiplexing/juggling classification, roughly split 60-40 between fast and slow juggling. This suggests that rapid switching within a trial was more prevalent than switching across trial. The remainder are relabeled overreaching under which the switching behavior is not tightly tied to the A and B spike count benchmarks. This further underscores the competitive strength of overreaching as an alternative, effectively addressing the gap in the previous framework. A minority of mixture triplets were classified as fixed-Poisson (Preferred, Non-Preferred, Middle), suggesting that SCAMPI model exhibited greater tolerance to overdispersion for the fixed Poisson category.

Among intermediate triplets, a majority were categorized as fixed-middle, indicating consistency between the two frameworks. A small subset of intermediate triplets was classified as fast-juggling, implying that in some cases information-preserving fluctuation may have occurred at a faster rate, a detail previously hidden within the intermediate category in the original model.

The remaining classifications under the original model mostly aligned with the results from the SCAMPI model. The majority of triplets labeled as single were classified as either fixed-preferred or fixed-non-preferred (with some of each). Additionally, all triplets labeled as outside were classified as fixed-outside(with some re-categorized as overreaching).

The disparity between the two frameworks offers evidence that the SCAMPI model enhances and refines the original model by enabling the detection of jugging within a trial (fast-juggling) and ruling out fluctuations unrelated to information preservation (overreaching).

### 4.5 Assessment of false discovery rate

Finally, to guard against false discoveries, we evaluated Bayesian False Discovery Rates (BFDRs) for each dataset: 33.5% (IC), 45.8% (IT-AL), 41.3% (IT-MF), 37.8% (V1-imp), 38.3%(V1-adj). BFDR represents the expected proportion of code juggling being false discoveries, calculated by averaging posterior probabilities of being non-code-juggling over the cases labeled as code juggling (fast-juggling and slow-juggling). The relatively high values of BFDR suggest that a non-negligible proportion of triplets are hard to accurately classify, leaving open the door to more precise statistical evaluation.

## 5 Discussion

This paper proposes new statistical tools for evaluating evidence of code juggling in neural activity under dual stimuli exposure relative to the activity patterns triggered by each constituent stimulus in isolation. Our findings suggest that although code juggling may not be detectable in every neuron in a population, it manifests within a non-negligible sub-population. This manifestation, in spite of its partial nature, is noteworthy because we believe code juggling could be a key computational tool for the brain to preserve multiple stimuli at little additional cost as the number of simultaneously presented stimuli increases. Let us elaborate.

On a first thought, it might seem unlikely that the brain should implement a flexible switching operation to preserve information about multiple stimuli. But the problem must be solved somehow and there are few compelling alternatives. It is unlikely that there is a lookup table with a unique value in the neural code for every possible stimulus combination that might exist. Such a computing strategy will be quickly defeated by the curse of dimensionality of the signal space. In contrast, the code juggling paradigm is not plagued by the curse of dimensionality of the signal space, as it takes advantage of task splitting over the temporal dimension so that every single neuron can alternate between encoding only a small set of signals that are within its receptive field. Such compositional computing strategies are massively scalable and could be efficient methods for preserving information from complex sensory scenes.

Although our analysis offers scientific evidence supporting the existence of code juggling in multiple brain areas, as well as its association to object identification, there is room for improvement. Indeed, our Bayesian false discovery rate analyses suggest that some of the triplet classifications are associated with low confidence. A partial explanation of this classification ambiguity is that spike counts mask a lot of the information about the neuronal activity dynamics that take place within the course of each trial. Consequently, spike count histograms could average over important features, diminishing the statistical separation between juggling and non-juggling activities. Future work will look into whether a deeper analysis of spike timing data could produce more precise statistical evaluation.

One potential extension is to incorporate over-dispersion in single-stimulus trials. Although neuronal response to a single stimulus is typically modeled as Poisson distributed around an expected count (Ventura et al., 2002; Kass et al., 2005), in some datasets many neurons fail to exhibit single-stimulus trials that conform to Poisson assumptions. Specifically, about half of neurons in the IC suffer from this problem, whereas in general more than 90% of neurons in visual areas survive screening for Poisson-like response patterns. To address this, we could extend the Poisson model to a negative binomial model and formulate corresponding biologically meaningful hypotheses, which may enable us to identify encoding modes for most recorded triplets. Another potential extension is to expand the dual-stimuli condition to multiple-stimuli conditions. However, developing a statistical framework that can accurately represent code juggling and other biologically meaningful modes under these more complex conditions may be challenging.

An additional avenue for extension could involve a more detailed analysis of non-benchmarked fluctuation. In the current framework, we introduce overreaching as a control for code juggling, encompassing notable cases where switches occur between mixture component means that fall outside the range defined by the benchmarks. Investigating the factors or mechanisms driving these deviations from benchmarks presents an intriguing research question. One possible explanation for this could involve attentional processes. During dual stimulus trials, attention might modulate the signal strength associated with each stimulus, potentially leading to shifts in the expected benchmark means. Another explanation could be the presence of subtle dynamics occurring on a finer timescale, which may be obscured by the trialwise aggregated spike counts. For example, each switch might incur a “cost” by briefly inhibiting neuronal firing. Although code juggling occurs within a trial, it may still be classified as overreaching because the mixture component means estimated from spike counts could deviate from the benchmark due to this inhibition. Future research could focus on conducting a more granular analysis of the underlying spike trains to investigate these phenomena further.

A broader issue that we can address in our future research is the neural coordination for information preservation. The code juggling patterns of a single neuron have paved the way for understanding how neural populations preserve information from multiple stimuli. According to a recent study by Jun et al. (2022), individual neurons exhibit fluctuating patterns that are positively and negatively related to other neurons with similar and different stimulus preferences, respectively. This finding demonstrates pairwise synchronous and asynchronous coordination among the neuron population. Moving forward, our future research will focus on detecting joint synchronous and asynchronous code-juggling, known as co-code-juggling, among multiple neurons. This investigation is an important step towards unraveling the mystery of the brain’s ability to perceive and represent complex sensory scenes.

## 6 Acknowledgements

We thank Marlene Cohen and Winrich Friewald for sharing their data and experimental details with us. The work has benefited from discussions with Cynthia King, Nicholas Marco, Meredith Schmehl, Justine Shih, Chad Smith, and Tingan Zhu. The work was partially supported NIH awards R01 DC013096 and R01 DC016363.

## Software

Coding for SCAMPI model is available in R package *neuromplex*. Related instructions on implementation are also provided.

## Supplementary Materials

### Figures and Tables

**Figure 8:**
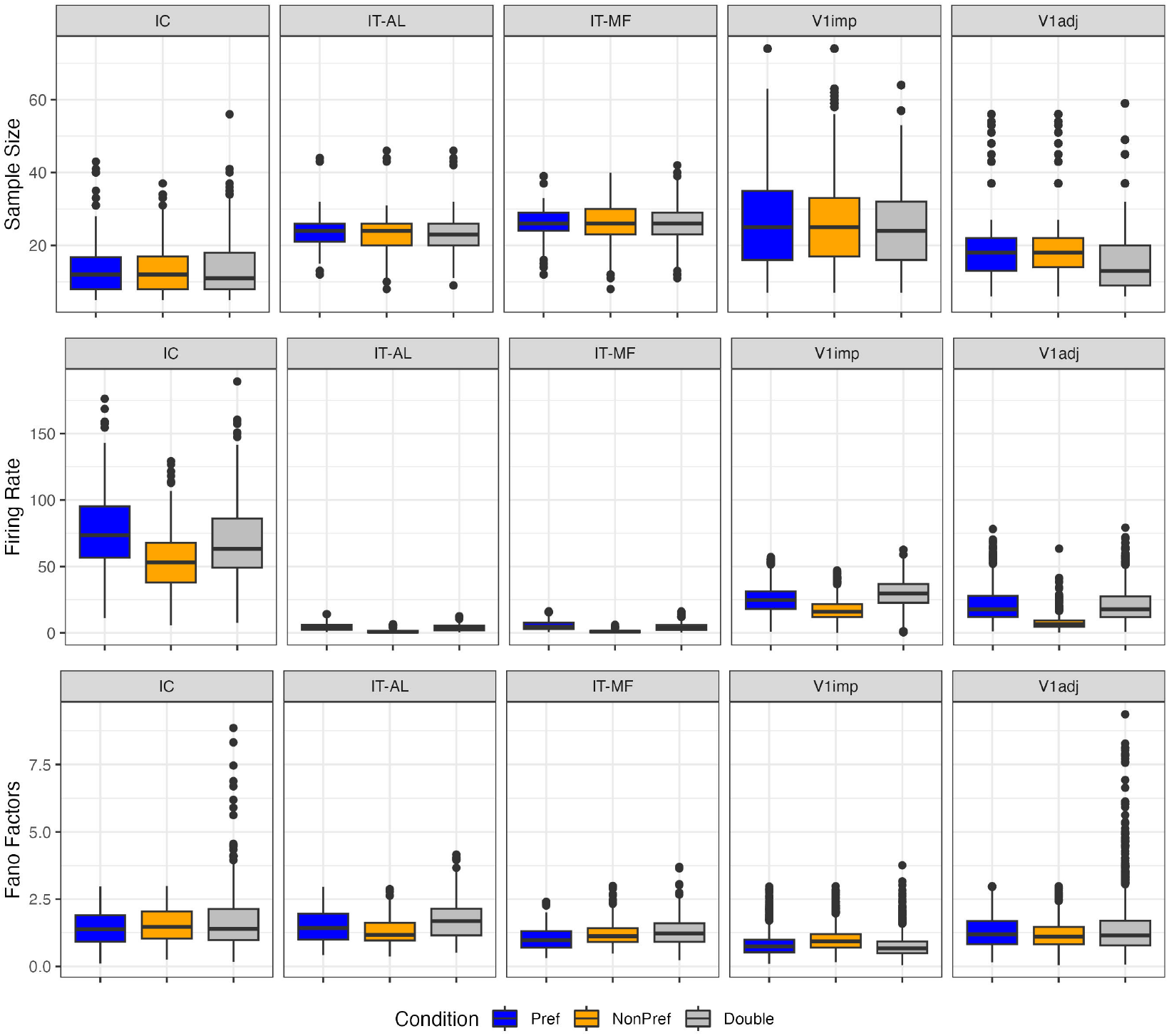
Summary statistics for each dataset.

**Figure 9:**
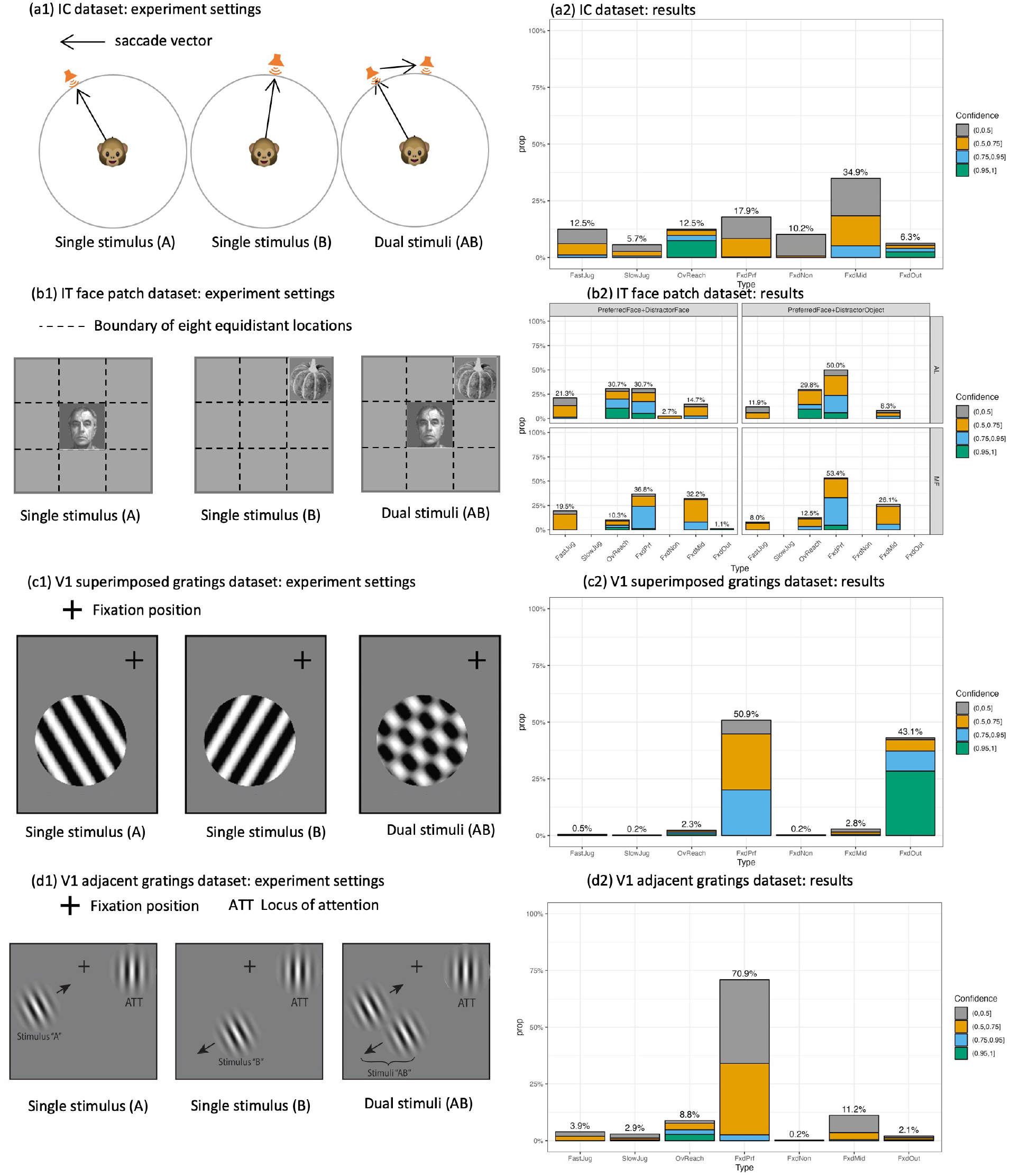
Summary of confidence levels for classification results for different datasets. Figures (a1)-(d1) present a selected experiment condition for a triplet defined in each of four datasets. Figures (a2)-(d2) visualize the corresponding classification results by the SCAMPI model, colored by posterior probability.

**Figure 10:**
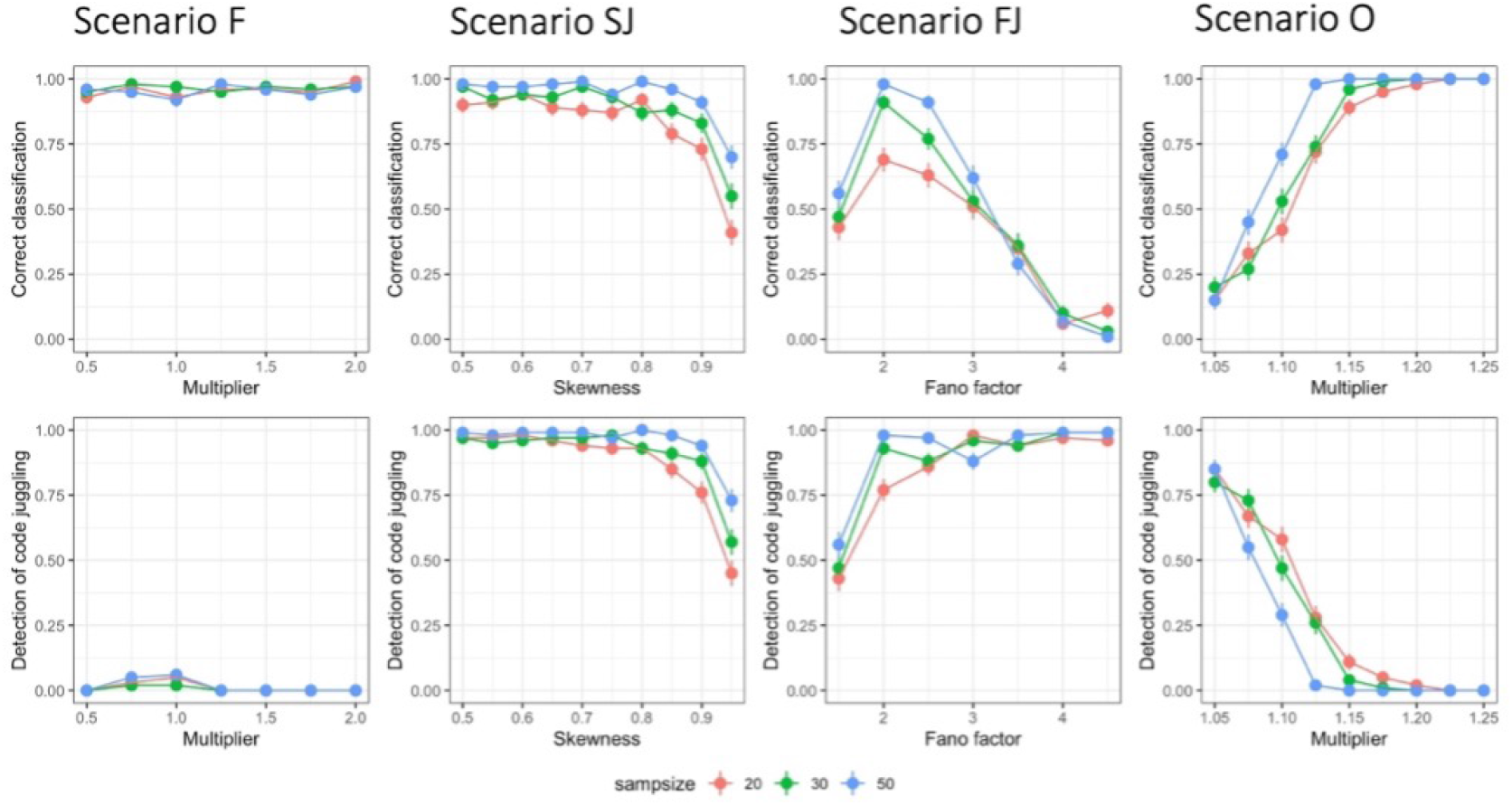
Performance evaluation of the SCAMPI model (frequentists’ two-stage framework). Classification accuracy and code juggling detection accuracy versus complexity parameters across fixed, slow-juggling, fast-juggling, and overreaching experimental cases (columns). Sample size 20, 30, 50 are color coded as red, green and blue.

**Figure 11:**
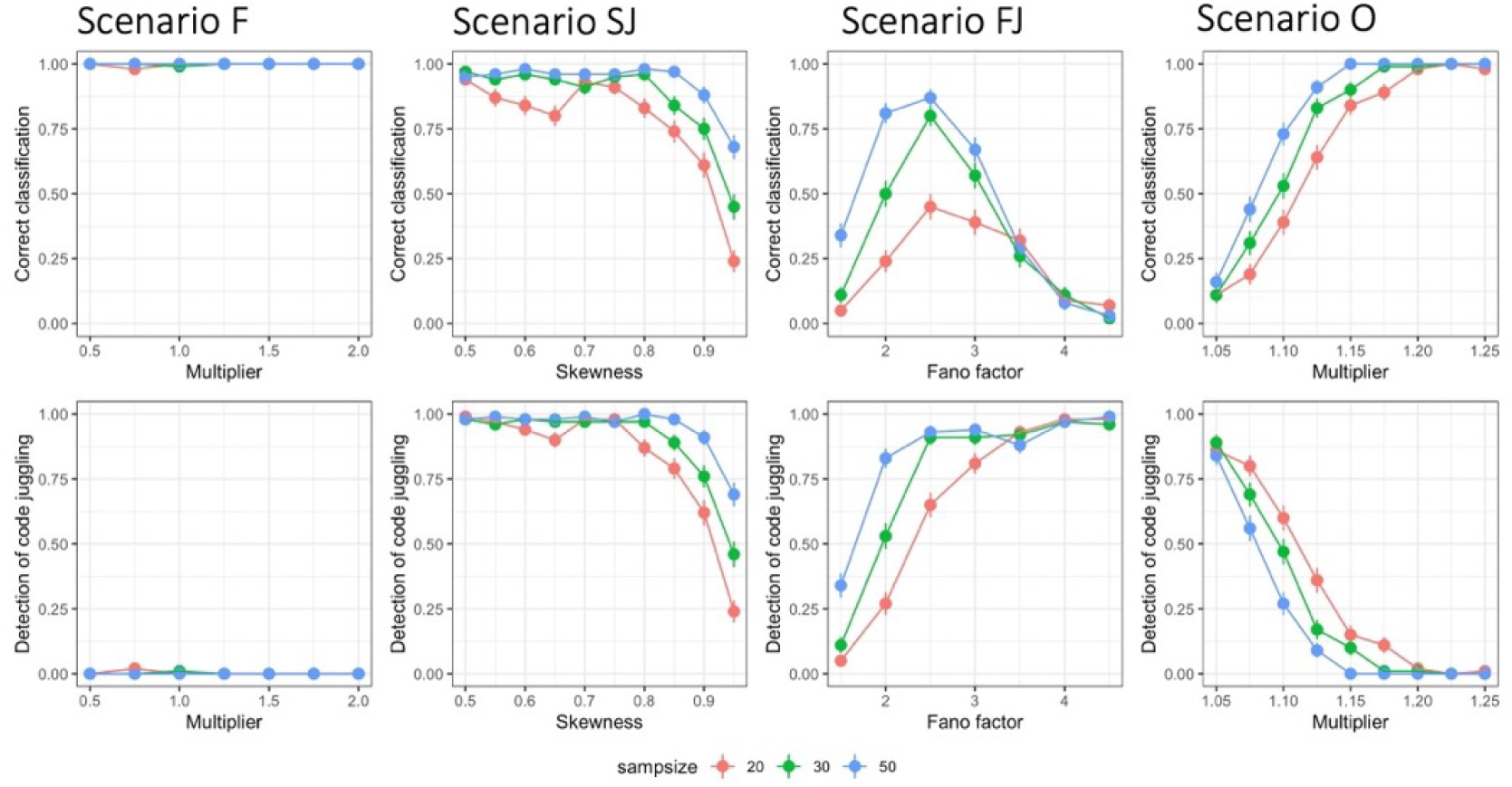
Performance evaluation of the SCAMPI model (intrinsic Bayes factor with Jeffreys’ prior for fixed hypothesis). Classification accuracy and code juggling detection accuracy versus complexity parameters across fixed, slow-juggling, fast-juggling, and overreaching experimental cases (columns).

**Figure 12:**
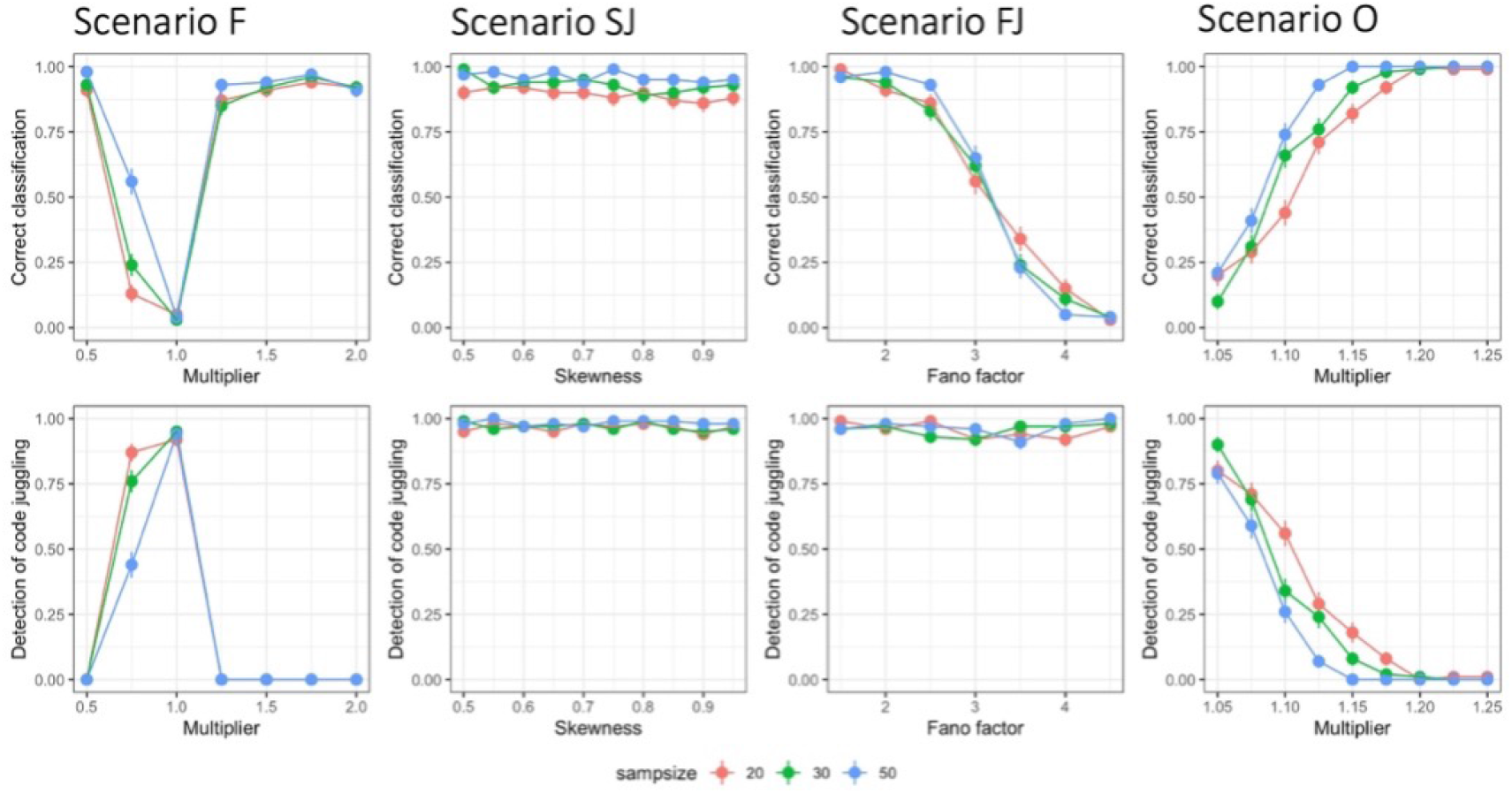
Evaluation of the SCAMPI model (bounded uniform prior for fixed hypothesis). Classification accuracy and code juggling detection accuracy versus difficulty tuning parameters across fixed, slow-juggling, fast-juggling, and overreaching experimental cases (columns).

**Figure 13:**
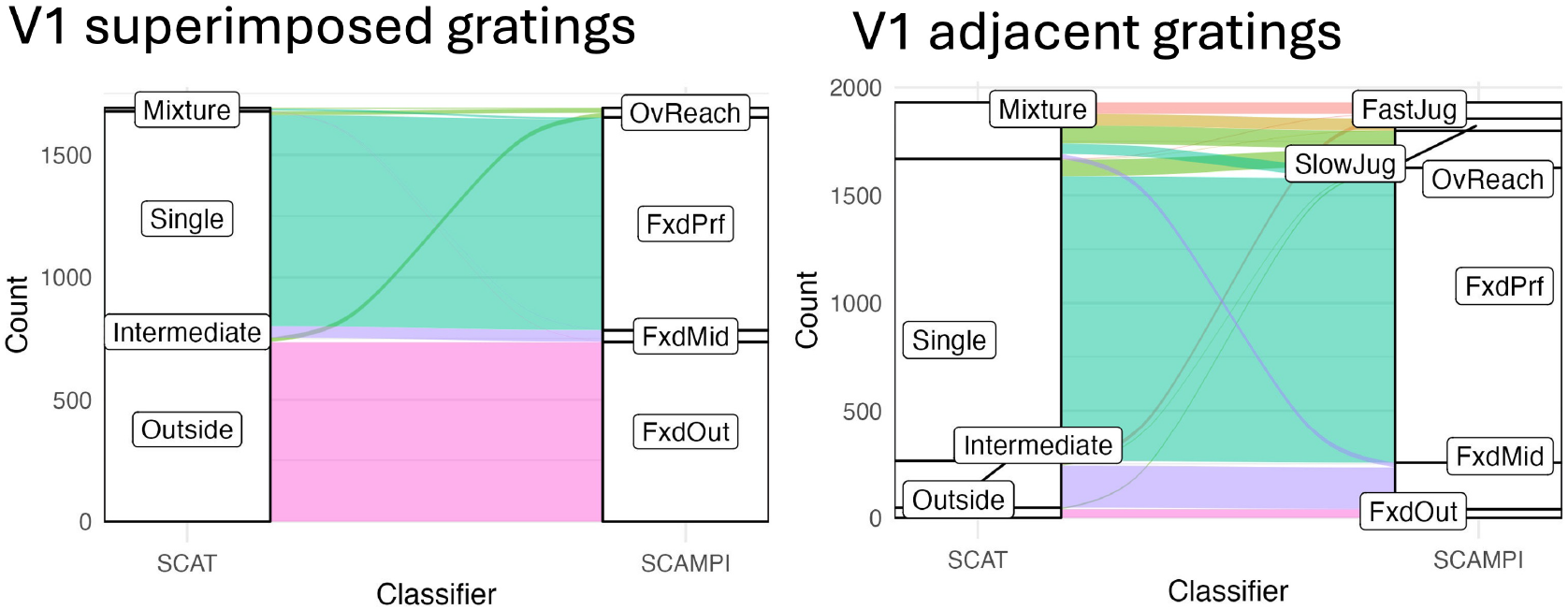
Results from both the original model and the SCAMPI model for the V1 dataset show the classifications under the SCAMPI model were largely consistent with the original model, especially when dual stimuli were superimposed gratings. For V1 adjacent gratings, disparities were most evident for mixture triplets, with equivalent proportions classified as fast-juggling, slow-juggling, and fixed-preferred. A slightly larger proportion of mixture triplets were classified as overreaching. Additionally, a very small proportion of single triplets were classified as overreaching.

**Figure 14:**
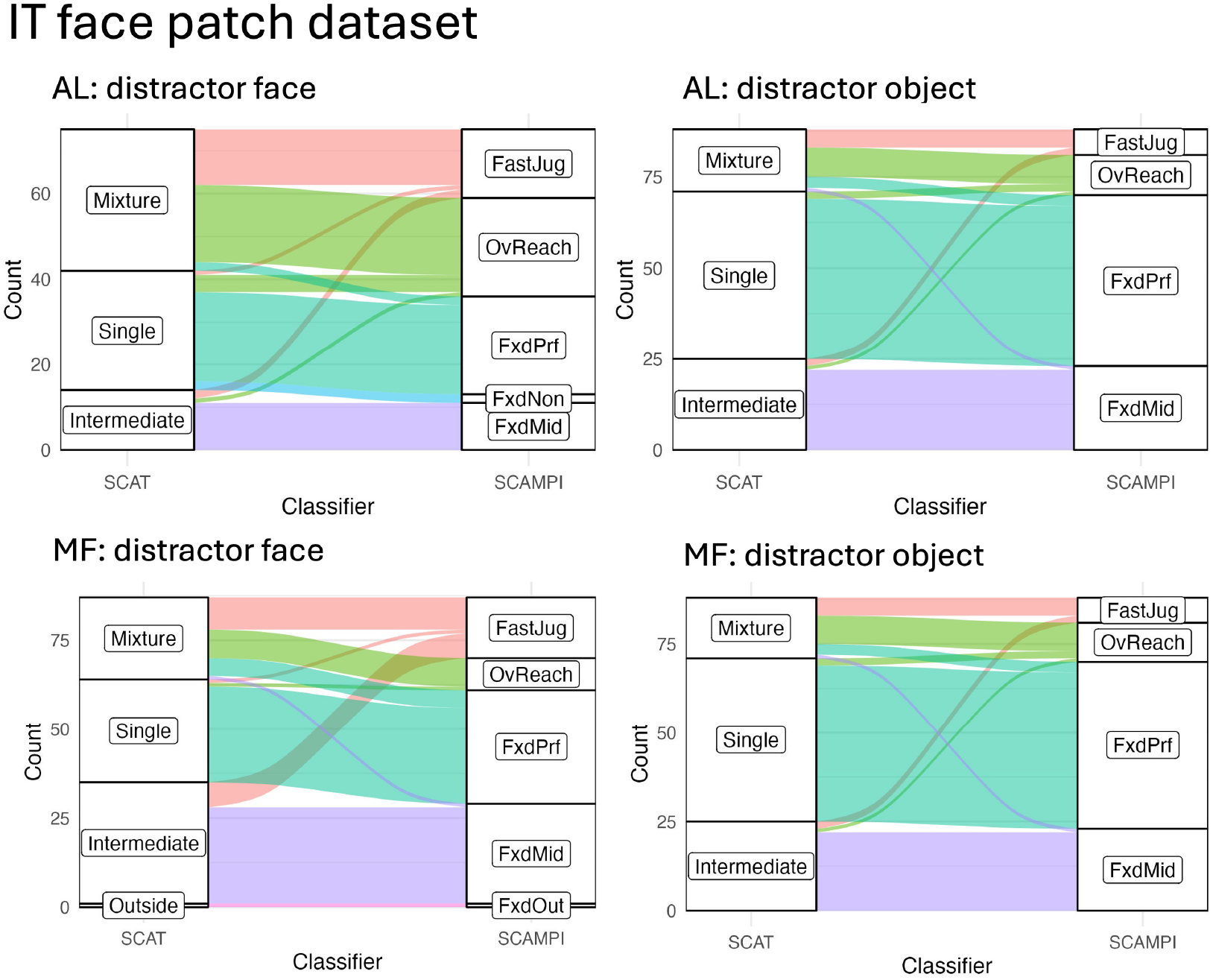
Results from both the original model and the SCAMPI model for the face patch dataset. Classifications under the SCAMPI model was largely consistent with the original model. mixture triplets were primarily classified as overreaching and fast-juggling. Additionally, a notable proportion of mixture triplets were classified as fixed-preferred. It is also noteworthy that a small proportion of intermediate triplets were classified as fast-juggling. However, neurons in the MF area with faces as distractors behaved differently compared to neurons in the AL area or when object were used as distractors. A larger proportion of intermediate triplets were classified as fast-juggling. The SCAMPI model provides a more refined classification of previously ambiguous cases.

### Technical Details

#### Gaussian Quadrature

For the integral calculation, we approximate the integral *∫*_Θ_· *dθ* with Gaussian quadrature. The basic idea is to approximate integral with a weighted sum of function values at specified points within the domain of integration. Here we set the points and weights according to Legendre polynomials proposed by Abramowitz and Stegun (1965).

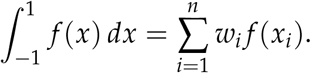

And change the interval from [−1, 1] to [*a, b*] according to

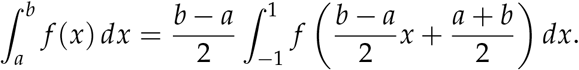

which is

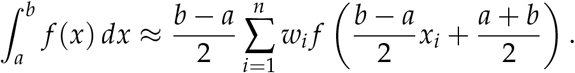

For two-dimension Gaussian quadrature, the calculation is similar. Here we also need to tranform the interval from [−1, 1] × [−1, 1] to [*a, b*] × [*c, d*]

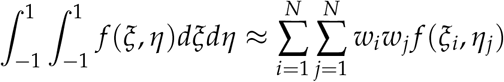

#### Laplace Approximation

According to Laplace approximation, approximate the posterior distribution with normal distribution.

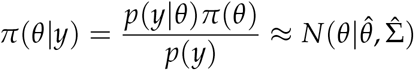

where

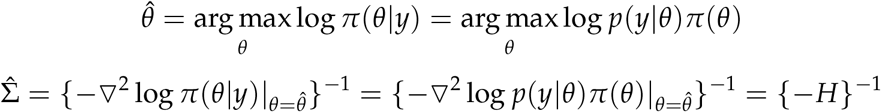

So we have the estimation for marginal likelihood valued at 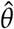:

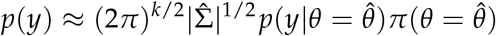

where *k* = *dim*(*θ*),*H* is Hessian matrix. So approximation on marginal likelihood is transformed to an optimization problem:

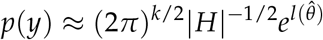

where 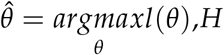,*H* is corresponding Hessian matrix.

#### Predictive Recursion Marginal Likelihood Gradient (PRMLG) Algorithm

##### Algorithm 1 Pseudocode for PRMLG Algorithm

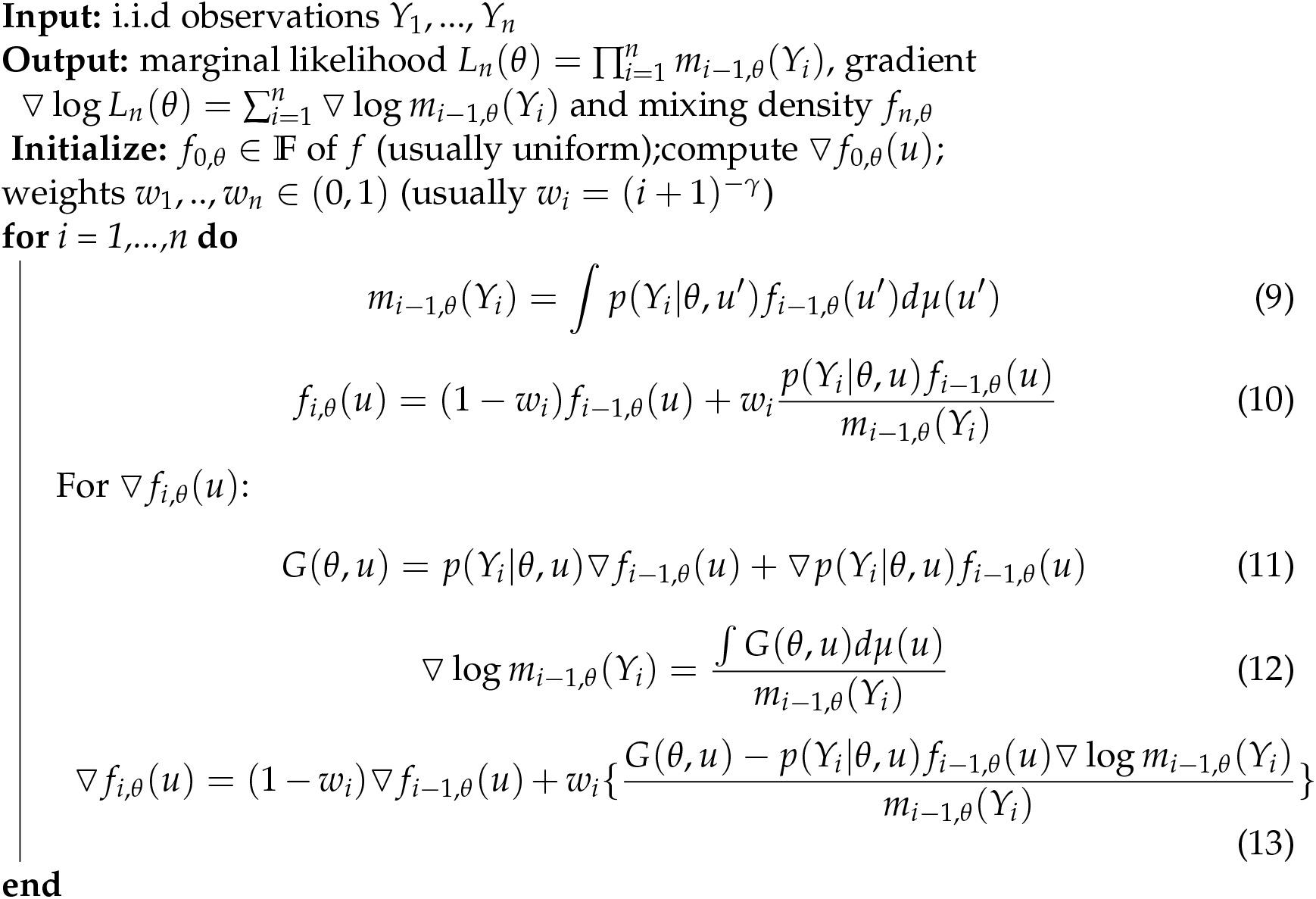

**Supplementary Details for Section 3: Fano Factor for** intermediate The data is denoted by *ϒ*, assumed to have a distribution *P* for intermediate, where

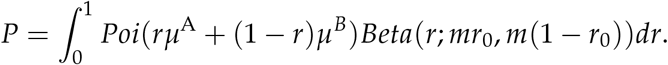

This can be rewritten as

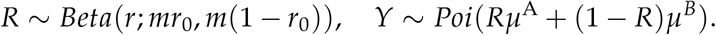

Denote the Fano factor of *ϒ* as *FF*(*ϒ*), we have

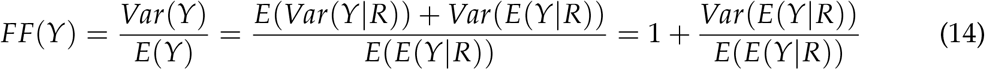

We can compute E(E(Y|R)) and *Var*(E(Y|R)) as follows:

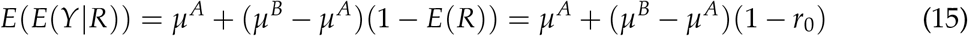

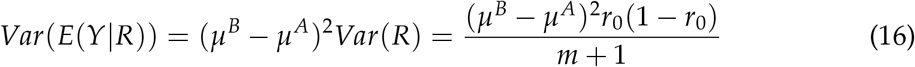

Plug equation (15) and (16) into equation (14), we have

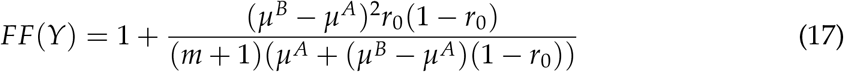

As shown in equation (17), *FF*(*ϒ*) monotonically decreases as *m* increases. When *m* approaches infinity, the Fano factor becomes 1. This corresponds to the case where fastjuggling degenerates into fixed, where *P* = Poi(*r*_0_*µ*^*A*^ + (1 − *r*_0_)*µ*^*B*^). In the scenario where *m* = 0, fast-juggling degenerates into slow-juggling, where *P* = *r*_0_Poi(*µ*^*A*^) +(1 − *r*_0_)Poi(*µ*^*B*^). In c this case, the Fano factor is a function of *r*_0_, and it reaches its maximum value when 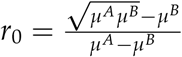.

In Section 3, we conducted simulations with *µ*^*A*^ = 50 and *µ*^*B*^ = 80. The maximum Fano factor of around 4.5 was achieved when *r*_0_ = 0.56 and *m* = 0, corresponding to the case where fast-juggling degenerates to slow-juggling *P* = 0.56Poi(50) + 0.44Poi(80). We kept *r*_0_ fixed at 0.56, and varied the precision parameter *m* from 0 to infinity, obtaining a wide range of Fano factors between 1 and 4.5. The minimum Fano factor of 1 was attained when *r*_0_ = 0.56 and *m* = 0, corresponding to the case where fast-juggling degenerates into fixed, with *P* = Poi(63.2).

## Footnotes

1 I.e., *P*^A^ < *P*^AB^ < *P*^B^ if *μ*^A^ < *μ*^B^ and *P*^B^ < *P*^AB^ < *P*^A^ otherwise, where *P*_1_ < *P*_2_ denotes stochastic ordering of probability distributions: *P*_1_((*x*, ∞)) ≤ *P*_2_((*x*, ∞)) for all *x*, with strict inequality for some *x*.

2 Under Poisson assumption for single-stimulus distributions *P*^A^, *P*^B^, testing separation between *P*^A^ and *P*^B^ is equivalent to test *μ*^A^ ≠ *μ*^B^ versus *μ*^A^ = *μ*^B^ = *μ*. Caruso et al. (2018)calculated the intrinsic Bayes factor sing equal prior probability assigned to two competing hypotheses and non-informative Jeffrey’s prior for unknown parameters. In our work, we followed Caruso et al. (2018)’s approach to retain the triplets with the logarithm of the intrinsic Bayes factor being equal or greater than three.

## Notes

### Competing Interest Statement

The authors have declared no competing interest.

## References

Abramowitz, M. and I. A. Stegun (1965). Handbook of mathematical functions: with formulas, graphs, and mathematical tables, Volume 55. Courier Corporation.

Berger, J. O. and L. R. Pericchi (1996). The intrinsic bayes factor for model selection and prediction. Journal of the American Statistical Association 91(433), 109–122.

Caruso, V. C., J. T. Mohl, C. Glynn, J. Lee, S. M. Willett, A. Zaman, A. F. Ebihara, R. Estrada, W. A. Freiwald, S. T. Tokdar, et al. (2018). Single neurons may encode simultaneous stimuli by switching between activity patterns. Nature communications 9(1), 2715.

Ebihara, A. F. (2015). Normalization among heterogeneous population confers stimulus discriminability on the macaque face patch neurons.

Festa, D., A. Aschner, A. Davila, A. Kohn, and R. Coen-Cagli (2021). Neuronal variability reflects probabilistic inference tuned to natural image statistics. Nature communications 12(1), 3635.

Glynn, C. D., S. T. Tokdar, A. Zaman, V. C. Caruso, J. T. Mohl, S. M. Willett, and J. M. Groh (2019). Analyzing second order stochasticity of neural spiking under stimuli-bundle exposure. Submitted.

Groh, J. M., M. N. Schmehl, V. C. Caruso, and S. T. Tokdar (2024). Signal switching may enhance processing power of the brain. Trends in Cognitive Sciences.

Jun, N. Y., D. A. Ruff, L. E. Kramer, B. Bowes, S. T. Tokdar, M. R. Cohen, and J. M. Groh (2022). Coordinated multiplexing of information about separate objects in visual cortex. Elife 11, e76452.

Kass, R. E., V. Ventura, and E. N. Brown (2005). Statistical issues in the analysis of neuronal data. Journal of neurophysiology 94(1), 8–25.

Martin, R. and S. T. Tokdar (2011). Semiparametric inference in mixture models with predictive recursion marginal likelihood. Biometrika 98(3), 567–582.

Mohl, J. T., V. C. Caruso, S. T. Tokdar, and J. M. Groh (2020). Sensitivity and specificity of a bayesian single trial analysis for time varying neural signals. Neurons, behavior, data analysis and theory 3(1).

Newton, M. A., F. A. Quintana, and Y. Zhang (1998). Nonparametric bayes methods using predictive updating. In Practical nonparametric and semiparametric Bayesian statistics, pp. 45–61. Springer.

Ruff, D. A., J. J. Alberts, and M. R. Cohen (2016). Relating normalization to neuronal populations across cortical areas. Journal of Neurophysiology 116(3), 1375–1386.

Ruff, D. A. and M. R. Cohen (2016). Attention increases spike count correlations between visual cortical areas. Journal of Neuroscience 36(28), 7523–7534.

Schmehl, M. N., V. C. Caruso, Y. Chen, N. Y. Jun, S. M. Willett, J. T. Mohl, D. A. Ruff, M. Cohen, A. F. Ebihara, W. A. Freiwald, et al. (2024). Multiple objects evoke fluctuating responses in several regions of the visual pathway. Elife 13, e91129.

Semedo, J. D., A. Zandvakili, C. K. Machens, M. Y. Byron, and A. Kohn (2019). Cortical areas interact through a communication subspace. Neuron 102(1), 249–259.

Tokdar, S. T., R. Martin, J. K. Ghosh, et al. (2009). Consistency of a recursive estimate of mixing distributions. The Annals of Statistics 37(5A), 2502–2522.

Ventura, V., R. Carta, R. E. Kass, S. N. Gettner, and C. R. Olson (2002). Statistical analysis of temporal evolution in single-neuron firing rates. Biostatistics 3(1), 1–20.

